# Folding-upon-binding pathways of an intrinsically disordered protein from a deep Markov state model

**DOI:** 10.1101/2023.07.21.550103

**Authors:** Thomas Sisk, Paul Robustelli

**Affiliations:** Dartmouth College, Department of Chemistry, Hanover, NH, 03755

**Author notes:** To whom correspondence should be addressed. Paul Robustelli: Address: 6128 Burke Laboratory Department of Chemistry Hanover, NH, 03755.

**Keywords:** Intrinsically Disordered Proteins, Molecular Recognition, Markov State Models, Deep Learning, Molecular Dynamics

## Abstract

A central challenge in the study of intrinsically disordered proteins is the characterization of the mechanisms by which they bind their physiological interaction partners. Here, we utilize a deep learning based Markov state modeling approach to characterize the folding-upon-binding pathways observed in a long-time scale molecular dynamics simulation of a disordered region of the measles virus nucleoprotein N_TAIL_ reversibly binding the X domain of the measles virus phosphoprotein complex. We find that folding-upon-binding predominantly occurs via two distinct encounter complexes that are differentiated by the binding orientation, helical content, and conformational heterogeneity of N_TAIL_. We do not, however, find evidence for the existence of canonical conformational selection or induced fit binding pathways. We observe four kinetically separated native-like bound states that interconvert on time scales of eighty to five hundred nanoseconds. These bound states share a core set of native intermolecular contacts and stable N_TAIL_ helices and are differentiated by a sequential formation of native and non-native contacts and additional helical turns. Our analyses provide an atomic resolution structural description of intermediate states in a folding-upon-binding pathway and elucidate the nature of the kinetic barriers between metastable states in a dynamic and heterogenous, or “fuzzy”, protein complex.

## Introduction

Intrinsically disordered proteins (IDPs) are proteins that do not adopt stable tertiary structures in isolation under physiological conditions. IDPs are ubiquitous in eukaryotic proteomes and viruses; and play crucial functional roles in many cellular processes.^1–3^ The biological functions of IDPs are often mediated by short sequence segments, referred to as linear motifs or molecular recognition elements, that interact with structured partner proteins.^4–6^ The molecular recognition elements of IDPs populate a structurally diverse set of conformations in their unbound states and can adopt a similarly diverse set of conformations when bound to different physiological interaction partners.^7–10^ This conformational plasticity enables IDPs to function as hubs in cellular signaling pathways, where they can form specific interactions with multiple binding partners.^11–13^ The relative affinities of these interactions can be tuned by post-translational modifications or changes in the cellular environment allowing for sensitive spatial and temporal regulation of cellular processes mediated by IDP interactions.^11, 14–18^

The thermodynamics of IDP interactions are complex, and the relationships between their free and bound state structures are not straightforward.^19^. In some instances, IDPs undergo disorder-to-order transitions and adopt stable tertiary structures when bound to physiological binding partners; a process referred to as “folding-upon-binding”.^5, 9, 20–22^ In other instances, IDPs retain a substantial amount of conformational disorder in their bound states.^23–26^ Such dynamic and heterogenous complexes are sometimes referred to as “fuzzy” complexes.^27, 28^ Substantial effort has been made to characterize the kinetics and thermodynamics of IDP binding events^6, 9, 29–31^, as elucidating the relationship between the free and bound states of IDPs will enable a more predictive understanding of their roles in biological pathways and human disease.^11, 32^

Stopped-flow and temperature-jump kinetics measurements^31, 33, 34^, NMR spectroscopy^35–39^, single molecule FRET^40–43^ and protein engineering techniques^44–46^ have emerged as powerful tools for characterizing the binding processes of IDPs. While these experimental techniques provide detailed mechanistic insight into IDP binding pathways, the data generated by these approaches are generally insufficient to obtain atomic resolution descriptions of the conformational states populated in IDP binding pathways. Atomistic descriptions of IDP binding intermediates and the conformational states populated by IDPs in complexes with their physiological interaction partners are highly desirable as they may facilitate the development of rational drug design strategies for modulating the activity of IDPs implicated in the pathogenesis of diseases.^17, 47, 48^

All-atom molecular dynamics (MD) computer simulations provide a powerful complement to biophysical experiments for characterizing conformational ensembles,^49–53^ binding pathways^44, 46, 54–56^ and bound states of IDPs.^48–53, 56–59^ Long timescale MD simulations run with an accurate physical model, or *force field*, can provide atomically detailed structural descriptions of conformational substates involved in IDP binding. MD simulations with sufficient statistical sampling of binding events also provide the equilibrium populations of these states and the rates of transitions between them.^54, 55^ Recent improvements to molecular mechanics force fields have dramatically enhanced the accuracy of MD simulations of disordered proteins and have shown promise for describing molecular recognition mechanisms of IDPs.^48, 52, 56, 58, 60, 61^ As IDP binding pathways occur on rugged and high-dimensional free energy surfaces, identifying mechanistically meaningful metastable states in MD simulations of IDP remains a substantial challenge.

Markov State Models (MSMs) describe the dynamics of stochastic systems as a transition network of memoryless, probabilistic jumps between sets of states. MSMs are a powerful approach for obtaining mechanistic insight from MD simulations^62, 63^ and have provided insights into protein conformational transitions^51, 64, 65^, protein folding^66^, protein-ligand binding^47, 55, 67^ and protein-protein complex formation.^47, 54, 55, 66–69^ The accuracy, interpretability, and relevance of information extracted from MSMs are, however, highly dependent on the input features used to describe a simulated system, the methods used to reduce the dimensionality of the input feature space and the partitioning of simulation frames into Markov states.^62, 70, 71^ These tasks are particularly challenging when building MSMs to describe the high-dimensional conformational space of disordered proteins.^47, 51, 72^

In recent years, theoretical advancements and applications of machine learning techniques have facilitated the construction of MSMs from MD simulation data.^73^ Automated feature selection, dimensionality reduction, and feature scoring methods can be applied to guide and validate the selection of molecular features to construct MSMs.^74–78^ These methods identify subsets of slowly evolving structural features, or *collective variables,* that can be used to partition MD trajectories into metastable Markov states that accurately model the kinetics of simulated conformational transitions.^76, 79, 80^ The variational approach to Markov processes (VAMP) has emerged as a powerful framework to identify molecular features that describe the slowest evolving degrees of freedom in a simulated system.^80–83^ In this approach a scoring function is used to quantify how effectively a set of features describes the kinetics of slow conformational transitions observed in MD simulations, and this score is maximized to identify optimal collective variables for MSM construction. The VAMP method has been extended to a deep learning framework where neural networks (referred to as “VAMPnets”) are optimized to identify metastable conformational states directly from molecular features.^84^ VAMPnet approaches have been further extended to include physical constraints in the training of neural networks that enable MSMs to be learned directly from simulation data.^85^ These models, referred to as “deep reversible MSMs”, “deep MSMs”, or “Koopman Models”, allow for the construction of kinetic models comprised of probabilistic states that may be differentiated by only subtle conformational features.^51, 85^

In this investigation, we have built a conventional MSM and a deep learning based MSM (or “deep MSM”) to characterize the folding-upon-binding pathways observed in a 200μs unbiased MD simulation of the α-helical molecular recognition element of the measles virus nucleoprotein N_TAIL_ reversibly binding the X domain (XD) of the measles virus phosphoprotein complex.^56^ The conformational dynamics of measles virus N_TAIL_ in solution and the folding-upon-binding of N_TAIL_ to XD have been extensively characterized by a variety of experimental^33, 36, 86–91^, and computational methods.^56, 92–94^ Here, we construct a hidden Markov state model^95^ using time-lagged independent component analysis (tICA)^79, 80, 96, 97^, a linear dimensionality reduction technique, and a deep MSM by applying the VAMPnet approach with physical constraints.^85^ Our deep MSM employs a multi-input neural network architecture that utilizes a combination of convolutional and fully connected neural network layers to merge structural descriptors with different inherent dimensionalities.

We find that the deep MSM identifies several states that were not identified by a conventional hidden Markov state model. The hidden Markov state model identifies a single heterogenous encounter complex state between N_TAIL_ and XD and a single heterogenous non-native complex where N_TAIL_ binds on the opposite face of XD relative native binding site. The deep MSM resolves two structurally and kinetically distinct encounter complex states that are differentiated by the binding orientation and helical content of N_TAIL_ as well as a kinetic trap on the native folding upon binding pathway. The deep MSM also identifies a network of several distinct non-native bound complexes. The hidden Markov state model and deep MSM both resolve 4 kinetically separated bound native-like states that interconvert on time scales of eighty to five hundred nanoseconds. These bound states share a core set of native intermolecular contacts and stable helices and are differentiated by a sequential formation of non-native contacts that facilitate the folding of additional helical turns. Interestingly, the detailed molecular mechanisms of folding-upon-binding revealed by our MSMs are not consistent with canonical conformational selection or induced-fit folding-upon-binding mechanisms. We find that encounter complexes that contain highly helical N_TAIL_ conformations proceed to the fully folded N_TAIL_:XD complex through a similar network of states as encounter complexes where N_TAIL_ has little helical structure.

Our analyses provide an atomic resolution structural and kinetic description of intermediate states in a folding-upon-binding pathway and elucidate the nature of the kinetic barriers between metastable states in a dynamic and heterogenous, or “fuzzy”, protein complex^10, 26–28, 98^ formed by an IDP and a structured binding partner. The neural network architecture designed here to train a deep MSM merges convolutional neural network layers that reduce the dimensionality of intermolecular contact matrices with fully connected network layers to describe global structural features. This neural network identifies several conformational states that were not resolved utilizing a reaction coordinate approach, time-lagged independent component analysis (tICA), or a conventional neural network architecture employing only fully connected neural network layers. These states enhance the resolution of the folding-upon-binding mechanism and suggest that folding-upon-binding proceeds through binding pathways that are inconsistent with canonical conformational selection or induced-fit binding mechanisms. This multi-input neural network approach may provide a general strategy for building deep MSMs to model the highly dynamic conformational states of IDPs and protein complexes with substantial conformational disorder.

## Results

### Molecular dynamics simulation of the measles virus nucleoprotein N_TAIL_ and the X domain of the measles virus phosphoprotein complex

A 200μs explicit solvent unbiased MD simulation of a 21-residue partially helical molecular recognition element of the measles virus nucleoprotein N_TAIL_ (residues 484-504, henceforth referred to as “N_TAIL_”) and the X domain (XD) of the measles virus phosphoprotein complex was previously performed by Robustelli et. al^56^ using the Anton^99^ supercomputer. This simulation was performed at 400 K using the a99SB-disp protein force field and a99SB-disp water model.^52^ A temperature of 400 K was selected for long time scale folding-upon-binding simulations as it was found to be near the simulated melting temperature of the N_TAIL_:XD complex and enabled an efficient sampling of binding and unbinding transitions in an equilibrium simulation. This simulation was initiated from an unbound conformation of N_TAIL_ and contains 36 binding and 36 unbinding events, where binding and unbinding events are defined using the fraction of native intermolecular contacts (*Q*)^56, 100^ as a reaction coordinate (See Methods). Here, we observed that XD unfolds at the beginning of this trajectory and refolds to its native state after 3 μs of simulation time and that XD unfolds and refolds multiple times in the final 30 μs of the trajectory. As we are only interested in modeling the binding pathways of N_TAIL_ to the native state of XD, we restricted our analysis to a continuous 167 μs subset of the original MD trajectory (from t=3 μs to t=170 μs) where XD remained in its native conformation. This 167 μs segment of the original trajectory contains 831701 frames, spaced with an interval of 200 ps per frame. We refer to this 167 μs segment as the “full trajectory”.

### Markov state model input features

We considered a set of input features containing 1029 intermolecular distances (one distance between each of the 21×49 intermolecular pairs of residues in N_TAIL_ and XD), 21 binary features based on the DSSP secondary structure assignment^101^ of each residue of N_TAIL_, and 15 features consisting of the value of the helical order parameter Sα^102^ for each consecutive seven residue fragment of N_TAIL_ (See Methods). We refer to sum of Sα values for all 15 seven residue fragments of N_TAIL_ as “N_TAIL_ Sα”. We consider a total of 1065 features for each MD simulation frame to build an 831701 x 1065 input feature matrix.

### Constructing a hidden Markov state model (HMSM) from time-lagged independent component analysis (tICA)

We utilized time-lagged independent component analysis (tICA)^79, 80, 96, 97^ to reduce the dimensionality of the N_TAIL_:XD input feature matrix and build an initial MSM. tICA was performed on the input feature matrix using a lag time of 6 ns and the resulting data were projected onto the first ten tICA eigenvectors. Initial analyses revealed that the binary DSSP assignment features had no impact on tICA projections and subsequent analyses, and they were subsequently excluded from the input features for building MSMs from tICA (See Methods). We visualize the free energy surface of the N_TAIL_:XD folding-upon-binding MD trajectory as a function of the two dominant time-lagged independent components (TICs) in Supplementary Figure 1. We observe that this projection resolves 4 distinct bound-state free energy basins that resemble the native N_TAIL_:XD complex observed by x-ray crystallography (PDB ID 1T6O)^86^. We determined an initial estimate of the optimal number of states for an MSM derived from the first ten tICA eigenvectors by iteratively applying the *k*-means algorithm with an increasing number of clusters until the resultant states no longer had statistically distinguishable properties in terms of the fraction of native intermolecular contacts (*Q*), Sα, radius of gyration (R_g_) and root mean squared deviation (RMSD) from the native complex. Using this approach, we found seven clusters to be optimal. We estimated a traditional MSM using these clusters as state definitions and a lag time of 24 ns. The implied timescales (ITS) of this model, however, were not converged or fully resolved. This MSM also failed to satisfy the generalized Chapman-Kolmogorov (CK) test^62^ (eq. 5), failing to reproduce the fastest processes observed in this system (data not shown).

To produce a valid model, we constructed an MSM with a larger numbers of initial states and coarse grained them to a smaller number states via the HMSM formulism introduced by Noe et al.^95^ We found that coarsening an initial twelve state MSM with seven resolved implied timescales (including the stationary process) to a seven state HMSM with a lag time of 6 ns yielded resolved and converged implied timescales and a valid CK-test (Supplementary Figure 2). We refer to this model as the “tICA HMSM”. We number these states HMSM state 1-7 in ascending order based on their similarity to the native complex, as assessed by the average values of the native intermolecular contact fraction (<*Q*>), N_TAIL_ Sα (<N_TAIL_ Sα >), R_g_ (<R_g_>) and RMSD from the crystal structure of the native complex calculated from all structures in each state (Supplementary Figure 3 and Supplementary Table 1). A network representation of the tICA HMSM with structural depictions of each state with the calculated mean first passage times (MFPTs) between them is displayed in Figure 1.

**Figure 1.**
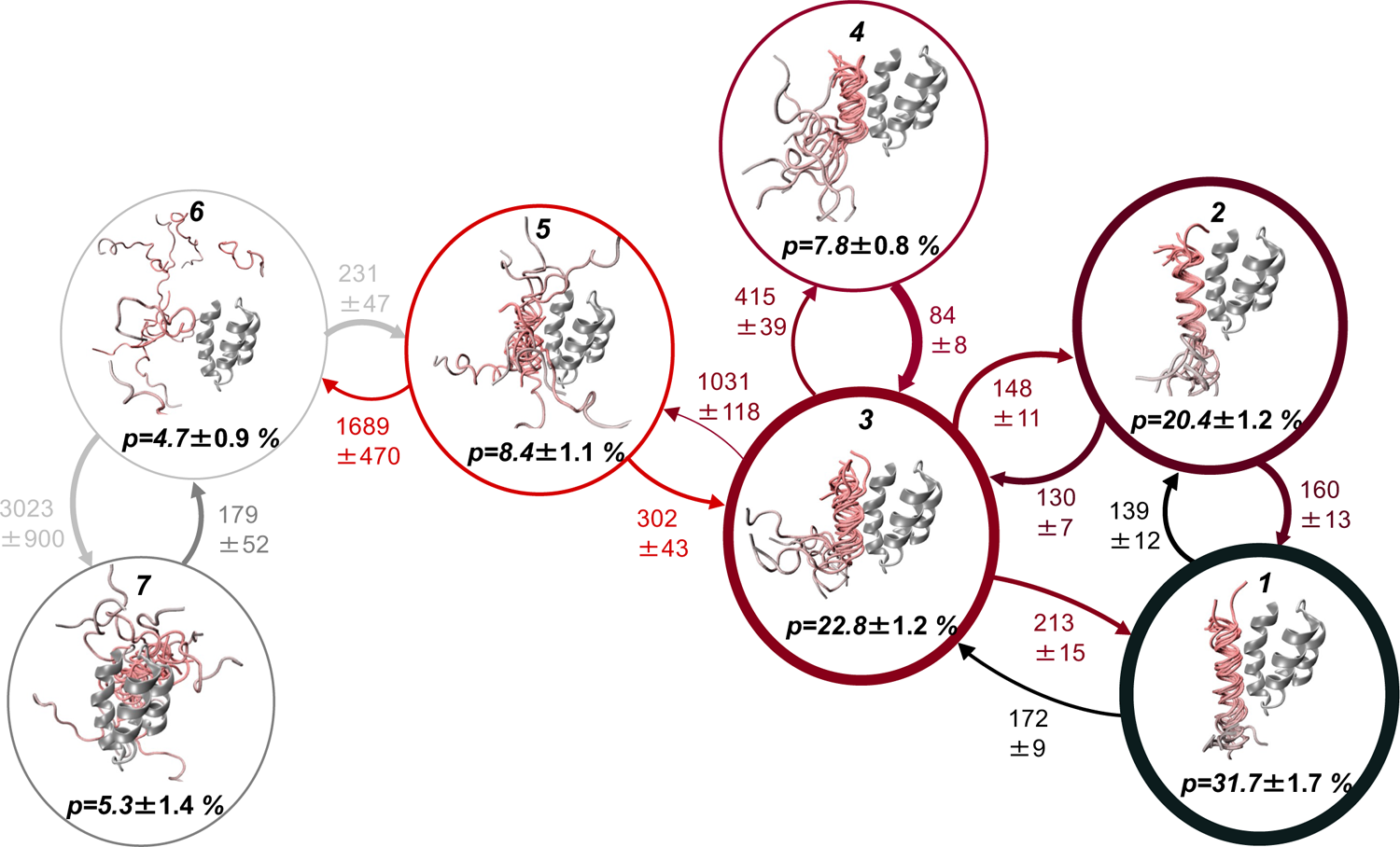
Transition network representation of a conventional hidden Markov state model of N_TAIL_:XD folding-upon-binding derived from a long-time scale equilibrium molecular dynamics simulation. Network representation of the transition matrix obtained from a hidden Markov state model (HMSM) derived from time-lagged independent component analysis (tICA) of a long-time scale MD simulation. Representative structures of each Markov state are displayed in circles along with their stationary probabilities (*p*). The thickness of circles is proportional to the stationary probability of each state. In representative structures of each state N_TAIL_ is colored with a gray-to-red gradient from the N-terminus to the C-terminus and XD is colored gray. Transition probabilities between states are indicated with directional arrows, and the thickness of the arrows is proportional the magnitude of the transition probability between states. Mean first passage times between states are reported in nanoseconds. All errors indicate the mean of the upper and lower deviations of the 95% confidence interval calculated from bootstrapping using 100 samples.

The HMSM state assignments are projected onto the two dominant tICs in Supplementary Figure 4. We visualize the free energy surface of each HMSM state as a function of the fraction of native intermolecular contacts (*Q*) and N_TAIL_ Sα in Supplementary Figure 5. The average values and standard deviations of *Q* and N_TAIL_ Sα for each HMSM state are compared in Supplementary Table 1 and Supplementary Figure 6. The populations of native and non-native N_TAIL_:XD intermolecular contacts and the N_TAIL_ helical propensities for each tICA HMSM state are compared in Supplementary Figure 7. The transition matrix of the HMSM is shown in Supplementary Figure 8 and the calculated MFPTs are shown in Supplementary Figure 9.

In HMSM state 1 N_TAIL_ adopts highly helical conformations (<N_TAIL_ Sα> = 10.9). These conformations have comparable helicity to the N_TAIL_ conformation observed in the native N_TAIL_:XD complex (N_TAIL_ Sα = 12.8 in PDB 1T6O) with the exception of helical fraying observed in the N-terminal N_TAIL_ residues G484-D487 and the C-terminal N_TAIL_ residues A502-I504. The average values of native intermolecular contacts <*Q*> are 0.93, 0.91, 0.79. and 0.78 and the average values of <N_TAIL_ Sα> are 10.9, 7.7, 5.6 and 4.8 for HMSM states 1-4, respectively. These 4 states contain stable helical conformations from N_TAIL_ A502 to A494 and are differentiated by the extension of stable N_TAIL_ helical conformations from N_TAIL_ A502 to D493, N_TAIL_ A502 to S491, and N_TAIL_ A502 to D487 in HMSM states 3, 2 and 1 respectively (Supplementary Figure 7). The R_g_ of the bound states increases from HMSM state 1 to HMSM state 4 as an increasing number of N-terminal residues of less helical conformations of N_TAIL_ extend outward from XD into solution (Supplementary Table 1). Our tICA HMSM also identifies a weakly bound state (HMSM state 5) with a small fraction of native intermolecular contacts and little helical content (<*Q*> = 0.16, <N_TAIL_ Sα> = 3.1), a state where N_TAIL_ and XD are largely unbound (HMSM state 6) with a substantially elevated R_g_ (<*Q*> = 0.01, <N_TAIL_ Sα> = 1.4, <R_g_> = 1.8 nm) and a more compact non-native complex (HMSM state 7) with very few native contacts (<*Q*> = 0.02, <N_TAIL_ Sα> = 3.6, <R_g_> = 1.3 nm) but more N_TAIL_ helical content than unbound N_TAIL_ conformations.

We observe that HMSM state 5 functions as a kinetic hub between unbound conformations in HMSM state 6 and the 4 native-like bound states (Figure 2, Supplementary Figure 8). HMSM State 5 can therefore be interpreted as an on-pathway encounter complex in the folding-upon-binding of pathway N_TAIL_. The most probable transitions from HMSM state 5 to the native-like bound states are to states 3 and 4, where N_TAIL_ is partially folded, with transition probabilities of 2.53 ± 0.4% and 1.48 ± 0.3%, respectively (error estimates computed with a Bayesian HMSM and Gibbs sampling approach^80, 112^, See Methods). From HMSM state 3, transitions to state 2 (7.33 ± 0.53%) are significantly more probable than to the less helical state 4 (4.85 ± 0.4%). HMSM state 4 has a relatively large probability of transitioning to state 3 (12.7 ± 1.03%) and very low probabilities of transitioning to states 1 (0.4 ± 0.1%) and 2 (1.1 ± 0.2%). Using Transition path theory (TPT)^103–105^, we find that folding-upon-binding pathways from HMSM state 6 (unbound) to states 1 and 2 (most native-like bound states) that exclude visits to state 4 comprise 74.8% of the total probability flux and that the pathway with the maximum flux (46.1%) proceeds through states 5, 3, and 2. We conclude that HMSM state 4 is largely off pathway to the more folded, bound states. We observe that HMSM state 7 consists of a non-native N_TAIL_:XD complex where N_TAIL_ is bound on the opposite face of XD relative to the native binding groove. Conformations in HMSM state 7 predominantly transition back to unbound conformations in HMSM state 6 (transition probability of 4.17 ± 0.78%) and very rarely transition directly to HMSM state 5 (transition probability of 0.1 ± 0.1%).

**Figure 2.**
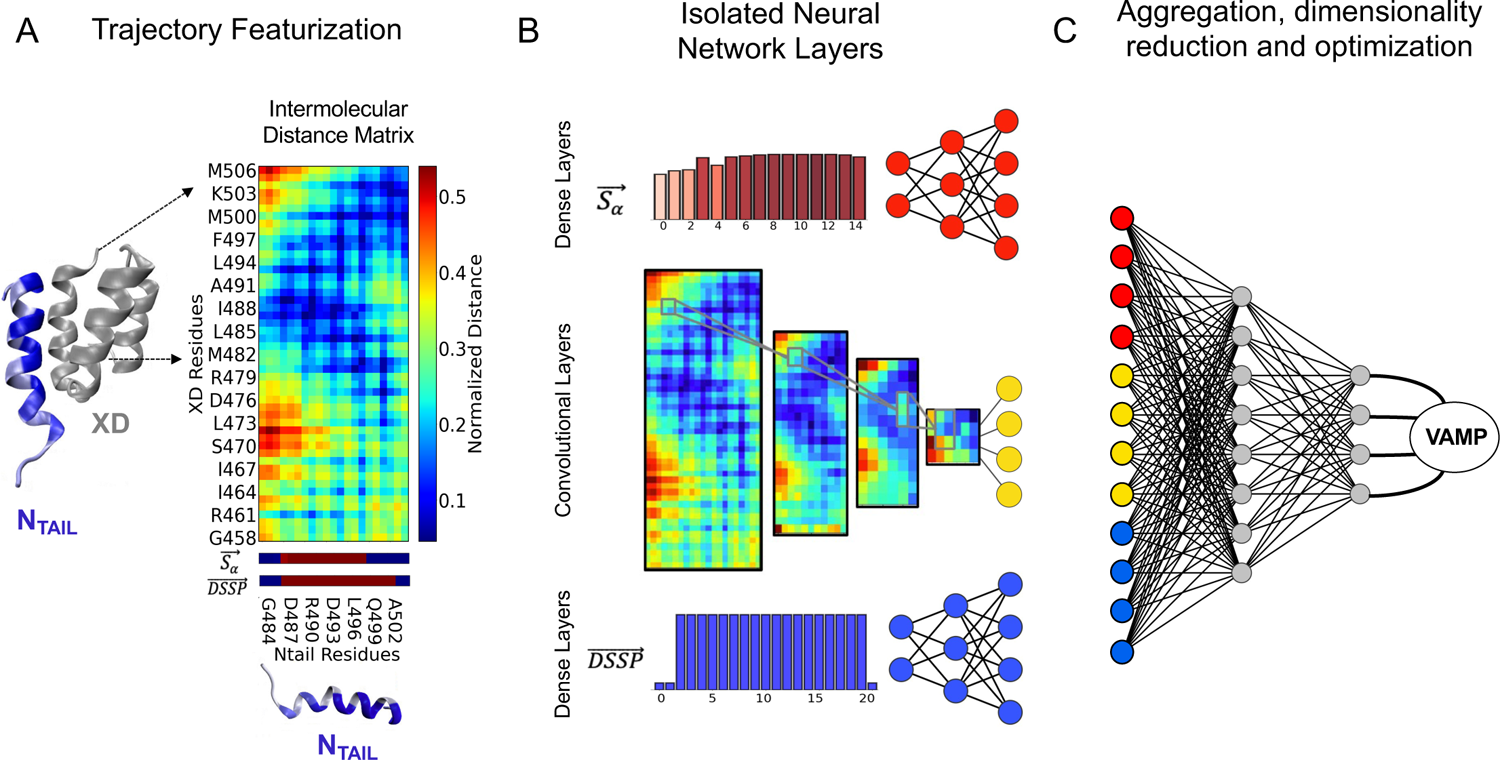
Multi-input neural network architecture used for building a deep Markov state model of N_TAIL_:XD folding-upon-binding. **(A)** Structural representation of the native N_TAIL_: XD complex. XD is colored gray and N_TAIL_ is colored in a gray-to-blue gradient proportional to each residues fraction of native contacts in deep MSM state 1. The set of deep MSM input features (intermolecular distances between N_TAIL_ and XD, N_TAIL_ Sα order parameters, binary DSSP helical assignments) are shown for the structure in A (right). **(B)** Schematic representation of the isolated neural network layers used to process each feature type based on its inherent dimensionality. Sα (red) and binary DSSP (blue) features are treated as 1D vectors and are processed with dense neural network layers. The intermolecular distance matrix between N_TAIL_ and XD is processed with convolutional neural network layers to take advantage of the spatial coherence of data points in its matrix form. **(C)** A qualitative schematic showing the aggregation and further processing of output features from the 3 isolated sets of layers. Upon aggregation, the processed output features from each isolated layer are combined by a final set of dense layers to reduce the dimensionality of the output to a normalized probability distribution over Markov states. The output probability distributions are used to compute a VAMP score for batches of time-lagged data pairs.

### Constructing a deep Markov state model with a multi-input neural network architecture

We sought to improve the resolution of our kinetic model and obtain greater mechanistic insight into N_TAIL_:XD folding-upon-binding by employing the deep learning “VAMPnet” approach with physical constraints to build a deep MSM.^85^ In this approach, the variational approach to Markov processes (VAMP) is integrated into a deep learning framework that combines feature selection, dimensionality reduction, state discretization, and kinetic modeling into a continuous pipeline for constructing MSMs. The VAMP provides a “VAMP score” that estimates how well a set of features describes the kinetics of the slowest evolving transitions observed in an MD simulation.^76, 81–83^ In a VAMPnet, a neural network is trained to learn a non-linear function that transforms input features into probabilistic state assignments that maximize the VAMP score. A VAMPnet outputs a probabilistic (or “fuzzy”) Markov state assignment for each frame of an MD simulation trajectory. Probabilistic state assignments describe the probability that each trajectory frame is a member of each Markov state. Higher VAMP scores result from probabilistic MSM state assignments that maximize the autocorrelation of each state assignment. Training neural networks to maximize VAMP scores therefore identifies slowly evolving state definitions describing metastable intermediates in long timescale processes.

Mardt et. al. extended the VAMPnet approach to learn a stochastic and reversible transition matrix defining the transition probabilities between fuzzy states obtained from an unconstrained VAMPnet.^84, 85^ A reversible and stochastic transition matrix adheres to detailed balance and has all positive elements so each element can therefore be interpreted as a transition probability. The learned deep MSM state assignments and reversible, stochastic transition matrix define a kinetic model from which the stationary distribution of states and their interconversion rates can be computed. These models have been referred to as “deep MSMs”, “VAMPnets with physical constraints” and “Koopman models” in previous studies due to their relationship with Koopman operator theory.^106^ Deep MSMs pose a great advantage over traditional MSMs as the utilization of neural networks in these models allow for the optimization of non-linear state membership functions.

We used the full set of 1065 input features to learn a deep MSM with a VAMPnet with physical constraints. We refer to this MSM as the “deep MSM”. To optimally integrate features that describe the helical content of N_TAIL_ (Sα and binary DSSP) and features that describe the position and orientation of N_TAIL_ relative to XD (the N_TAIL_:XD intermolecular distance matrix) in our VAMPnet, we designed a multi-input neural network architecture. A schematic illustration of this multi-input neural network architecture is presented in Figure 2. This neural network architecture employs a combination of convolutional network layers and fully connected network layers to merge structural descriptors with different dimensionalities. Convolutional neural networks provide dramatic performance advantages for deep learning tasks involving image data.^107^ Recognizing that the intermolecular distances matrix (or intermolecular “contact map”) between N_TAIL_ and XD obtained in each frame of the simulation can be interpreted as an image, we sought to leverage the local spatial coherence in these contact maps by transforming them with convolutional neural network layers in our VAMPnet. We then combine the information obtained from convolutional neural network layer transformations of intermolecular contact maps with information obtained from fully connected dense neural network layer transformations of the Sα and binary DSSP helical assignment features.

The three neural network inputs (intermolecular distance matrices, N_TAIL_ Sα values and binary DSSP N_TAIL_ helical assignments) are transformed separately in three branches, applying convolutional neural network layers to transform intermolecular contact maps and fully connected neural network layers to transform the vector quantities of Sα and binary DSSP helix assignments (See Methods). The resulting outputs from each branch of the network are combined and transformed by a final set of fully connected neural network layers. The details of the final architecture of this neural network are described and illustrated in Supplementary Figure 10. The initial fully connected neural network layers used to transform Sα values and binary helical DSSP assignments increase the dimensionality of these data to better capture relationships between different sequence regions in N_TAIL_ and the initial convolutional network layers reduce the dimensionality of intermolecular contact maps to better capture essential relationships between intermolecular contacts in different regions of the N_TAIL_:XD complex with a coarser representation of intermolecular distances.

We determined the final architecture of our neural network implementation and VAMPnet hyperparameters (batch size, learning rate, epsilon parameter, model lag time, and number of states) by iteratively optimizing the VAMP2 score (eq. 8) of an unconstrained neural network (See Methods). We found that using 12 output states and a lag time of 2 ns to train unconstrained VAMPnets maximized the VAMP2 score and consistently produced the same set of 12 distinguishable states. We characterize the latent space and state assignments of the initial unconstrained VAMPnet in Figure 3.

**Figure 3.**
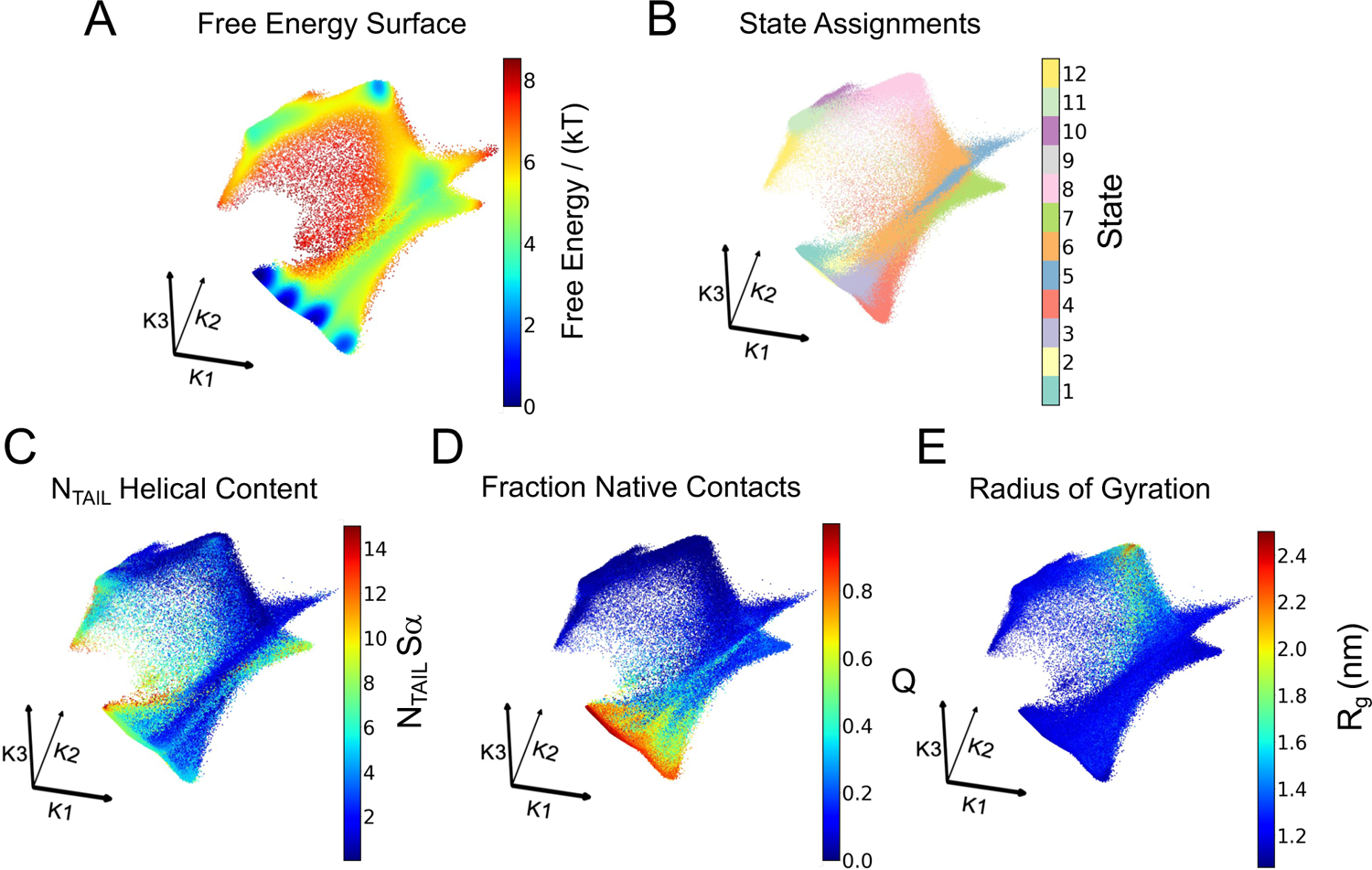
VAMPnet latent space and state assignments used to construct a deep Markov state model of N_TAIL_:XD folding-upon-binding. We characterize the latent space of our N_TAIL_:XD VAMPnet by projecting MD observables onto the left singular functions (or “Koopman modes”) K1, K2, and K3 of the half-weighted Koopman matrix estimated from an initial unconstrained VAMPnet. Truncating the singular value decomposition to 3 singular vectors gives a 3-dimensional latent space or set of singular functions where points are embedded in a kinetically meaningful way. We characterize the latent space representation of each MD simulation frame by coloring each data point by **(A)** the apparent free energy obtained by taking the negative natural log of a gaussian kernel density estimate over the 3-dimensional latent space-projected data, **(B)** its crisp Markov state assignment **(C)** the fraction of native intermolecular contacts (Q) **(D)** the sum of the N_TAIL_ α-helical folding order parameter Sα for each 7 residue segment of N_TAIL_ and **(E)** the radius of gyration (R_g_) of all Cα carbons of N_TAIL_ and XD.

We constructed our final deep MSM by retraining the initial unconstrained VAMPnet with physical constraints to learn a reversible and stochastic transition matrix defining the transition probabilities between the 12 states identified by the unconstrained VAMPnet (See Methods).^84, 85^ The Chapman-Kolmogrov (CK) test^62^, implied timescales, and steady state distributions for the deep MSM estimated at a lag time of 6 ns are shown in Supplementary Figure 11. We refer to the 12 states obtained from the deep MSM as deep MSM states 1-12. We number the states of the deep MSM in ascending order based on their similarity to the native N_TAIL_:XD complex in terms of the fraction of native intermolecular contacts (*Q*), N_TAIL_ Sα, radius of gyration and RMSD from the native complex (Supplementary Figure 12). We visualize the free energy surface of each deep MSM state as a function of *Q* and N_TAIL_ Sα in Supplementary Figure 13. We compare the average values and standard deviations of *Q*, N_TAIL_ Sα and the radius of gyration for each deep MSM state in Supplementary Table 2 and Supplementary Figure 14. We compare the populations of native and non-native N_TAIL_:XD intermolecular contacts and the N_TAIL_ helical propensities for each deep MSM state in Supplementary Figure 15. The transition matrix and the mean first passage times for the deep MSM are shown in Supplementary Figures 16 and 17, respectively.

A transition network representation of the deep MSM with structural depictions of each state and the mean first passage times between states is displayed in Figure 4. We observe that 5 of the deep MSM states closely resemble 5 of the tICA HMSM states. Deep MSM states 1-4 closely resemble the 4 native-like HMSM bound states (HMSM states 1-4). Deep MSM state 8, where N_TAIL_ is unbound, closely resembles HMSM state 6. In the tICA HMSM, we resolve a single heterogenous encounter complex state (HMSM state 5). The deep MSM increases the resolution of our model and effectively fine grains this heterogenous encounter complex into 3 distinct states: deep MSM states 5, 6 and 7. These states are substantially more homogenous than HMSM state 5 and are differentiated by the helical content of N_TAIL_, the orientation of N_TAIL_ relative to XD, the conformational heterogeneity of N_TAIL_ and the populations of native and non-native intermolecular contacts (Figures 4-5, Supplementary Figures 12-15).

**Figure 4.**
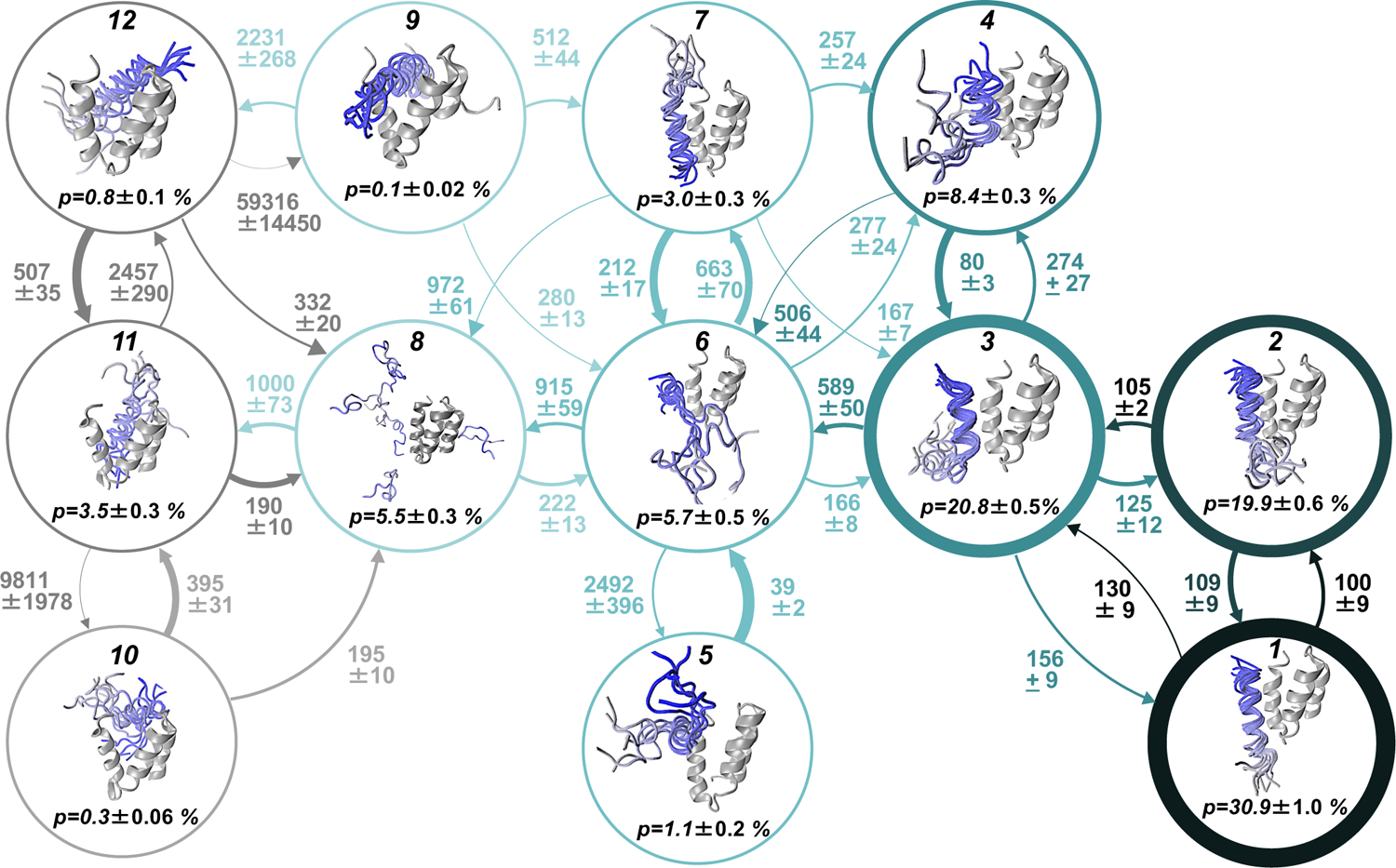
Transition network representation of a deep Markov state model of N_TAIL_:XD folding-upon-binding derived from a long-time scale molecular dynamics simulation using a multi-input neural network architecture. Network representation of the transition matrix of a deep Markov state model (MSM) obtained from a multi-input neural network architecture. Representative structures of each Markov state are displayed in circles along with their stationary probabilities (*p*). The thickness of circles is proportional to the stationary probability of each state. In the representative structures of each state, N_TAIL_ is colored by a gray-to-blue gradient from the N-terminus to the C-terminus and XD is colored gray. The transition probability between states is indicated with directional arrows, and the thickness of the arrows is proportional the magnitude of the transition probability between states. Mean first passage times between states are reported in nanoseconds. The values and errors reported here are the bootstrap means and their 95% confidence intervals, obtained from 30 independent optimization runs of the constrained VAMPnet.

**Figure 5.**
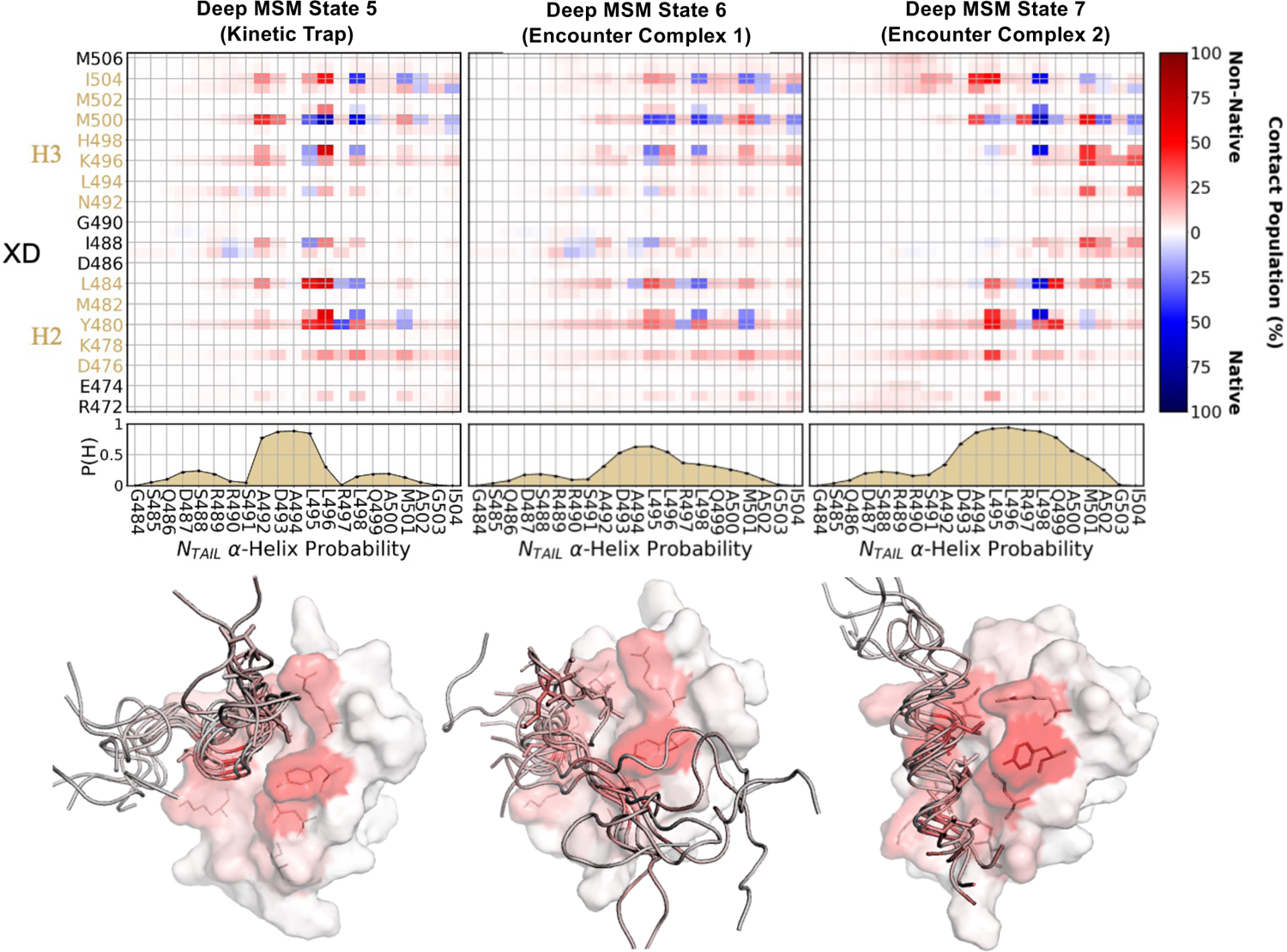
A deep Markov state model of N_TAIL_:XD folding-upon-binding pathways resolves two distinct encounter complex states and a kinetic trap state. State averaged intermolecular contact populations and N_TAIL_ helical propensities for deep MSM states 5, 6 and 7. Intermolecular contacts are defined as occurring in all frames where the minimum distance between heavy atoms of two residues is less than 5.0 Å. Native intermolecular contacts are colored blue and non-native contacts are colored red. Native contacts are defined as those present in the crystal structure (PDB 1T6O) using the same criteria. Helical propensities (P(H)) were calculated using DSSP. Structural representations contain an overlay of multiple representative N_TAIL_ structures with one surface representation of XD. The residues of N_TAIL_ and XD are colored by a gray-to-red gradient that represents the fraction of frames where non-native intermolecular contacts are formed by each residue in each state.

The deep MSM similarly fine grains HMSM state 7, the heterogenous non-native complex where N_TAIL_ is bound to the opposite face of XD relative to the native binding site, into 3 distinct states (deep MSM states 10-12, Figure 5, Supplementary Figures 12-15). In these more homogenous states N_TAIL_ is bound in different locations on XD and contains distinct populations of helical content. In addition, the VAMPnet also identifies a rare conformational state (deep MSM state 9, steady-state population *p* = 0.1 ± 0.02%) in which N_TAIL_ is inserted between the three helical bundles of XD.

### A deep Markov state model resolves two structurally and kinetically distinct encounter complex states and a kinetic trap

In the tICA HMSM, most of the probability flux from unbound N_TAIL_ states to native-like bound states flows through a single Markov state (tICA HMSM state 5) which functions as an encounter complex and kinetic hub for transitions between bound and unbound conformations (Figure 2, Supplementary Figures 8-9). tICA HMSM state 5 has a steady-state population (*p*) of *p* = 8.7 ± 1.1% and contains a small fraction of native intermolecular contacts (<*Q*> = 0.22) and relatively little helical content (<N_TAIL_ Sα> = 3.1). In the deep MSM this state has effectively been split into three states: deep MSM states 5, 6, and 7 (Figures 4-5). Deep MSM states 5, 6 and 7 have steady state populations of *p* = 1.1 ± 0.2%, *p* = 5.7 ± 0.5% and *p* = 3.0 ± 0.3%, respectively. We observe that the populations of helical N_TAIL_ conformations are substantially smaller in deep MSM state 5 (<N_TAIL_ Sα> = 1.4) and deep MSM state 6 (<N_TAIL_ Sα> = 2.0) compared to deep MSM state 7 (<N_TAIL_ Sα> = 5.1). We find that deep MSM states 5, 6 and 7 have similar fractions of native intermolecular contacts (<*Q*> = 0.19, <*Q*> = 0.19, and <*Q*> = 0.18, respectively) but observe that there is a large difference in the subsets of the intermolecular residue pairs that form native and non-native intermolecular contacts in each state (Figure 5).

N_TAIL_ residues L495 and L498 insert into the hydrophobic binding groove of XD in the native complex. In deep MSM state 6 these leucine residues form similar populations of native and non-native intermolecular contacts and N_TAIL_ is not restricted to native-like binding orientations, and instead samples a relatively isotropic distribution of rotational orientations. In deep MSM state 7, native intermolecular contacts formed by N_TAIL_ L498 have substantially higher populations than native intermolecular contacts formed by N_TAIL_ L495, and N_TAIL_ L495 forms highly populated non-native intermolecular contacts. Visual inspection of deep MSM state 6 and state 7 reveals that N_TAIL_ L498 binds at similar positions in the native XD hydrophobic binding groove in both states (Figure 5). In deep MSM state 7, however, N_TAIL_ L495 is inserted into a non-native binding site in the hydrophobic binding groove of XD that orients N_TAIL_ in the opposite (or “upside-down”) orientation relative to the N_TAIL_ orientation observed in the native N_TAIL_:XD bound complex. We define a rotational order parameter in the form of an angle to quantify the orientation of N_TAIL_ relative to the native binding face of XD in each deep MSM state in Supplementary Appendix 1 and present the distribution of this order parameter for each deep MSM state in Supplementary Figure 18.

N_TAIL_ conformations in deep MSM state 6 have a similar helical propensity to unbound states of N_TAIL,_ except for a slightly elevated helical propensity observed in residues A492-L495 (Figure 5, Supplementary Figure 15). In deep MSM state 7, N_TAIL_ has a higher helical propensity that more closely resembles the less helical native-like bound states (deep MSM states 3 and 4). One might therefore hypothesize that deep MSM state 6 functions as an encounter complex for a binding pathway resembling an “induced fit” mechanism, where the formation of native intermolecular contacts proceeds the subsequent folding of secondary structure elements formed the bound state, while deep MSM state 7 functions as an encounter complex for a parallel binding pathway resembling a “conformational selection” mechanism, where preformed native-like secondary structure elements bind XD before the subsequent formation of native intermolecular contacts. A detailed inspection of the transition probabilities and transition rates among deep MSM states, however, reveals that N_TAIL_ binding pathways do not fall into such a dichotomy (Figure 4, Supplementary Figures 16-17).

While N_TAIL_ conformations in deep MSM state 7 are substantially more helical than N_TAIL_ conformations in state 6, we do not observe greater transition probabilities from state 7 to the more helical native-like bound states 1 and 2 (Figure 5, Supplementary Figure 16). The transition probabilities from deep MSM state 7 to states 1 and 2 are 0.1 ± 0.01% and 0.5 ± 0.1%, respectively. These values are smaller than the transition probabilities observed from the less helical encounter complex (deep MSM state 6) to states 1 and 2 (0.6 ± 0.1% and 1.7 ± 0.2%, respectively). The highest transition probabilities from deep MSM state 7 are to state 6 (15.3 ± 0.8%) and state 4 (4.9 ± 0.7%), states where N_TAIL_ is substantially less helical.

These observations contrast with the classical paradigm of conformational selection, where a stable, preformed helix binds and remains helical for the duration of a binding event. We observe that the transition rates from the two deep MSM encounter complex states (states 6 and 7) to the deep MSM native-like bound states (states 1-4) are within statistical error (Supplementary Figure 17) and that deep MSM states 6 and 7 are most clearly kinetically distinguished based on incoming transitions from unbound and non-native conformations (Supplementary Figure 16). These results indicate that while we identify distinct encounter complex states with different N_TAIL_ helical propensities and conformational pathways leading to their formation, these states ultimately transition to native-like bound states with similar rates and ultimately form the same network of partially bound and folded fuzzy complexes that subsequently transition to the most native-like state. Consequently, we conclude that folding-upon-binding pathways originating from these encounter complex states are not well described by an induced fit / conformational selection dichotomy.

Deep MSM state 5 transitions almost exclusively to state 6 which is the only state that has an appreciable probability of transitioning to state 5 (Supplementary Figure 16). Consequently, we identify deep MSM state 5 as an off-pathway kinetic trap on folding-upon-binding pathways that proceed through state 6. Deep MSM state 5 is similar to state 6 but N_TAIL_ has an elevated helical propensity in residues A492-L495 (Figure 5). Deep MSM state 5 contains more highly populated non-native contacts between N_TAIL_ residues L495 and L496 and XD residues Y480, L481, L484, F497, and I504 (average population of 55.7 ± 7.33%) than state 6 and state 7 (average populations of 19.7 + 2.4% and 12.3 ± 4.7%, respectively). We thus identify the stabilization of helical conformations of N_TAIL_ by the formation of non-native contacts as the basis for the substantial kinetic barrier observed between deep MSM state 5 and the native-like bound states.

### Kinetic barriers between native-like bound states originate from non-native contacts

N_TAIL_ folding-upon-binding pathways from encounter complex states (deep MSM states 6 and 7) to the nost native-like bound state (deep MSM state 1) are largely mediated by the sequential formation and subsequent breakage of two distinct sets of non-native intermolecular contacts. The majority of the probability flux from the deep MSM encounter complex states to the native state states proceeds through deep MSM states 3 and 4. These states contain similar N_TAIL_ helical propensites and populations of native intermolecular contacts, but are differentiated by an elevated population of a cluster of non-native intermolecular contacts between N_TAIL_ residues A492 and D493 and XD residues D487, I488, and D493 in deep MSM state 3 (Figure 6). This cluster of non-native intermolecular contacts is highlighted by a dotted rectangle in Figure 6A, and representative depictions of these contacts are shown in Figure 6B. The average population of the non-native contacts between these groups of residues is 32.9 ± 0.7% in deep MSM state 3 compared to 3.9 ± 3.8% in state 4.

**Figure 6.**
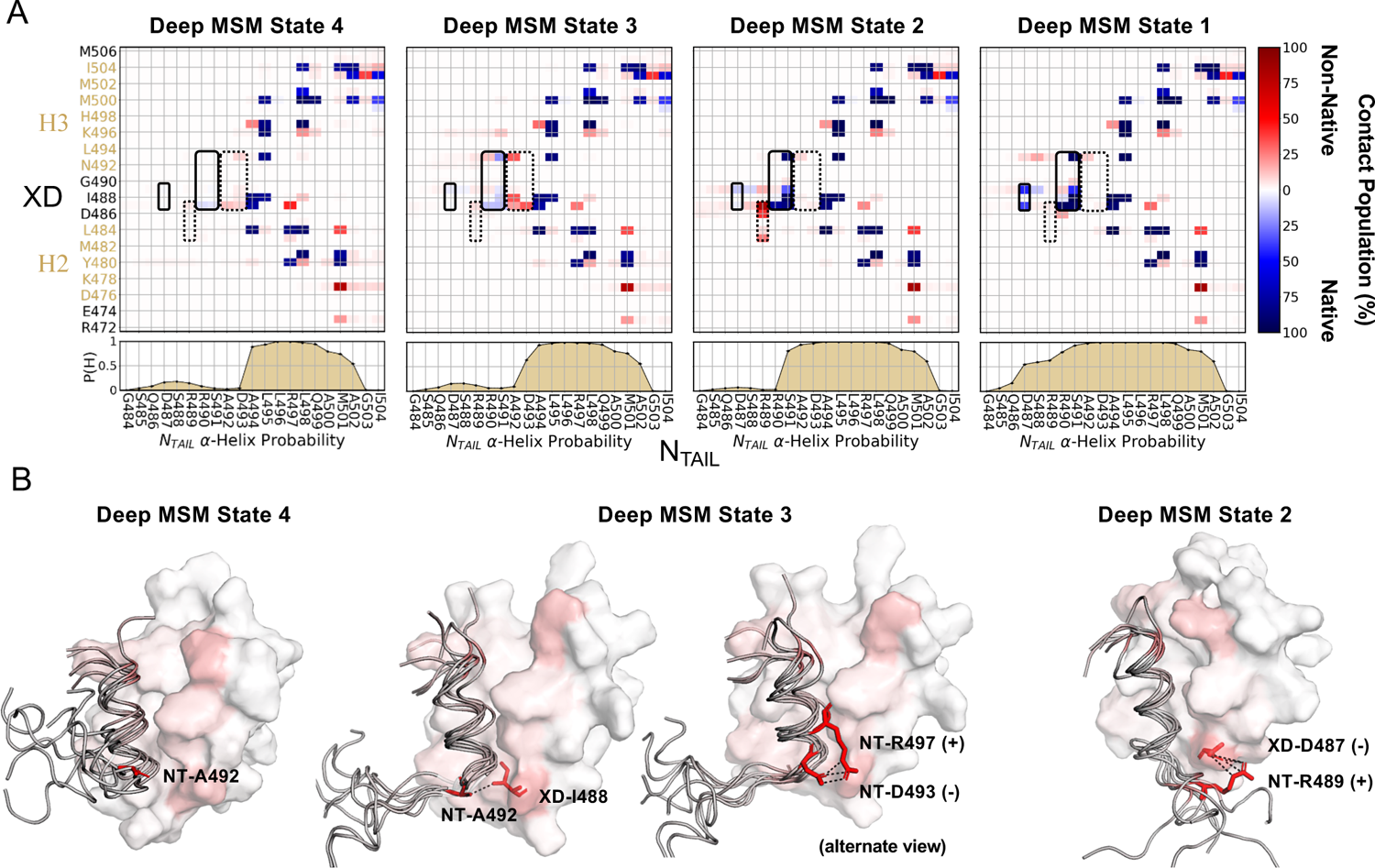
Kinetic barriers between native-like N_TAIL_:XD bound states originate from non-native intermolecular and intramolecular contacts. **(A)** State averaged intermolecular N_TAIL_:XD contact populations and N_TAIL_ helical propensities for native-like N_TAIL_:XD bound states. Intermolecular contacts are defined as occurring in all frames where the minimum distance between heavy atoms of two residues is less than 5.0 Å. Native intermolecular contact pairs are colored blue and non-native intermolecular contact pairs are colored red. Native contacts are defined as those present in the crystal structure (PDB 1T6O) using the same criteria. **(B)** Structural representations of native-like N_TAIL_:XD bound states. Each state representation is an overlay of multiple representative N_TAIL_ structures with one surface representation of XD. The residues of N_TAIL_ and XD are colored by a gray-to-red gradient that represents the fraction of frames where non-native intermolecular contacts are formed by each residue in each state. Selected sidechains of N_TAIL_ (NT) and XD are shown as sticks to illustrate important non-native contacts in different states.

Deep MSM state 3 contains a substantially populated intramolecular salt bridge between residues N_TAIL_ R497 and N_TAIL_ D493. We define this salt bridge as being formed in trajectory frames where one of the carbonyl oxygens of N_TAIL_ D493 is within 3.5 Å of a guanidinium nitrogen of N_TAIL_ R497. By this definition, the N_TAIL_ R497:D493 salt bridge has a population of 4.1 ± 0.7% in state 4 and 26.6 ± 0.1% in state 3. These results suggests that the kinetic barrier between deep MSM state 4 and state 3 partially results from the process of forming and breaking the intramolecular N_TAIL_ R497:D493 salt bridge and non-native intermolecular contacts between N_TAIL_ A492 and D493 and XD residues D487, I488, and D493. We observe that the process of forming these contacts is substantially faster than the process of breaking them (MFPT = 80.0 ± 3.4 ns for transitions from deep MSM state 4 to state 3 and MFPT = 274.1 ± 27.4 ns for transitions from state 3 to state 4). Interestingly, it has been observed that the N_TAIL_ mutation R497G substantially diminishes the affinity of N_TAIL_ to XD.^108^ K_D_ values of 3.0 ± 0.2 μM and 44.4 ± 2.2 μM were measured for wild type and R497G N_TAIL_, respectively. N_TAIL_ R497 forms stable native intermolecular contacts with XD in all the deep MSM native-like bound states. The absence of these native intermolecular interactions should destabilize the native complex between the N_TAIL_ R497G mutant and XD. The absence of an intramolecular salt bridge between N_TAIL_ R497 and D493 may further destabilize deep MSM state 3. As most of the total probability flux (70.7 ± 6.0%) from the unbound state (deep MSM state 8) to most native-like bound (state 1) proceeds through state 3, this additional destabilization of state 3 may contribute to the dramatic affinity loss observed for N_TAIL_ R497G observed in previous studies.

The formation of non-native intermolecular contacts in deep MSM state 3 coincides with the transient formation of several weakly populated native intermolecular contacts between N_TAIL_ residues R490 and S491 with XD residues D487, I488, and D493 (average population of 14.0 ± 4.4%, dark rectangle, Figure 6B). These native contacts subsequently become “locked in” after transitions to deep MSM state 2, where they have an average population of 86.7 ± 5.4%. The formation of these stable native intermolecular contacts is accompanied by a substantial increase in the population of intermolecular hydrogen bonds between the sidechain hydroxyl hydrogen of N_TAIL_ S491 and the carboxylic acid oxygens of XD D493 and the hydroxyl oxygen of N_TAIL_ S491 and the backbone amide hydrogen of XD K489. These hydrogen bonds are observed in the x-ray structure of the N_TAIL_:XD complex^86^ and the N_TAIL_ mutation S491L was previously demonstrated to reduce the affinity of N_TAIL_ to XD beneath the detection limits of ITC^108^, underscoring the importance of these intermolecular hydrogen bonds in stabilizing the N_TAIL_:XD complex. These hydrogen bonds have a population of 53.0 ± 0.4% in deep MSM state 2 compared to in 6.6 ± 0.1% and 0.2 ± 0.1% of frames in states 3 and 4, respectively. The formation of this cluster of native contacts in deep MSM state 2 is accompanied by an increase in the helical propensities of N_TAIL_ residues S491-D493, and the formation of several non-native intermolecular contacts between N_TAIL_ residue R489 and XD residues T483, D486 and D487 (average population = 49.3 ± 24.3%). The strongest non-native intermolecular contacts in this cluster occur between N_TAIL_ R489 and XD D487 (*p* = 82.5 ± 0.76%) and N_TAIL_ R489 and XD D486 (*p* = 40.2 ± 0.4%), demonstrating the importance of non-native intermolecular salt bridge interactions in stabilizing this state.

The stability of non-native contacts formed by N_TAIL_ R489 and XD residues T483, D486 and D487 appear to substantially contribute to the kinetic barrier between deep MSM state 2 and state 1. These contacts have an average population of 49.3 ± 24.3% in deep MSM state 2 but are nearly absent in state 1 (average population = 2.0 ± 1.7%). Transitions from deep MSM state 2 to state 1 are also accompanied by the formation of stable helical conformations from N_TAIL_ S491 to D487 and the formation of a final set of native intermolecular contacts between N_TAIL_ D487 and XD D487 and N_TAIL_ D467 and XD K489 (*p* = 37.7 ± 0.3% and *p* = 43.2 ± 0.3% respectively in deep MSM state 1). These native intermolecular contacts are indicated by a solid block box in Figure 4A. Transitions between deep MSM state 2 and state 1 are relatively fast (MFPT = 109.1 ± 7.2 ns for transitions from state 2 to state 1 and MFPT = 99.8 ± 10.7 ns for transitions from state 1 to state 2) and are among the fastest of the transitions observed between native-like bound states. This transition involves the cooperative extension of the N_TAIL_ helix by 4 residues, whereas the helix of N_TAIL_ is extended by only a single residue in transitions from deep MSM state 4 to state 3. The transition from deep MSM state 2 to state 1 involves the formation of a favorable salt bridge between N_TAIL_ D487 and XD K489 in a conformation where the aliphatic residues of N_TAIL_ D487 and XD D487 sidechains are in contact, but the negatively charged carboxylic acid moieties are orientated to minimize unfavorable charge interactions. We speculate that the strong electrostatic attractions and repulsions between this set of charged sidechains may facilitate the relatively fast transitions observed between deep MSM state 2 and state 1.

### Comparison of Markov state models with a 1D reaction coordinate for folding-upon-binding

In a previous investigation by Robustelli et. al^56^ a 1D reaction coordinate was optimized to characterize the folding-upon-binding mechanism observed in the MD simulation analyzed here. This reaction coordinate was derived using the fraction of native intermolecular contacts (*Q*) between N_TAIL_ and XD as an initial reaction coordinate and employing the variational optimization approach of Best and Hummer^109^ to reweight the contribution of each native intermolecular contact to produce a new reaction coordinate (*R*). This optimization was carried out to increase the maximum value of the conditional probability distribution p(TP|*R*), where p(TP|*R*) is the probability that a frame of the MD trajectory is on transition path at a given value of the optimized reaction coordinate *R*.

A projection of the MD trajectory onto the previously calculated 1D reaction coordinate *R* was found to contain three apparent free-energy minima separating unbound and native-like bound conformations (Supplementary Figure 19). It is, however, unclear if the apparent free-energy barriers observed in this projection are kinetically meaningful. We have calculated the probability distribution of the value of the reaction coordinate *R* for each kinetically distinct deep MSM state in Supplementary Figure 19. We observe that the two primary encounter complex states identified in this investigation (deep MSM states 6 and 7) are largely indistinguishable based on this reaction coordinate. We also observe that native-like bound states of the deep MSM (deep MSM states 1-4) are similarly indistinguishable based on this reaction coordinate. This result is unsurprising given the importance of non-native contacts in differentiating the Markov states of our deep MSM and underscores the complementary insights that MSMs can provide to low dimensional reaction coordinate approaches for describing protein folding and disordered protein folding-upon-binding.

## Discussion

We report the construction of Markov state models (MSMs) to structurally and kinetically characterize folding-upon-pathways observed in an unbiased long time scale MD simulation of a disordered molecular recognition element of the measles virus nucleoprotein N_TAIL_ reversibly binding the X domain of the measles virus phosphoprotein complex. We constructed a hidden Markov state model (HMSM) using time-lagged independent component analysis (tICA), a linear dimensionality reduction technique, and a deep learning based MSM (or “deep MSM”) using the VAMPnet approach with physical constraints with a multi-input neural network architecture. The MSMs constructed with these two approaches both resolve an unbound state and 4 kinetically separated native-like bound states that interconvert on time scales of eighty to five hundred nanoseconds. In the HMSM built using tICA, we observe that transitions between unbound N_TAIL_ conformations and native-like bound states of N_TAIL_:XD complexes predominantly occur through a single conformationally heterogenous Markov state, which we refer to as an “encounter complex” state. In contrast, the deep MSM built using the reversible VAMPnet approach resolves several additional structurally and kinetically distinct states including two encounter complexes and an off-pathway kinetic trap.

In both encounter complex states identified in the deep MSM N_TAIL_ residue L498 is inserted into the hydrophobic binding groove of XD in its native binding site. These encounter complex states are differentiated by the binding orientation, helical content, and conformational heterogeneity of N_TAIL_. In one encounter complex state N_TAIL_ adopts relatively disordered conformations with similar helical content to unbound N_TAIL_ conformations and samples a relatively isotropic distribution of rotational orientations relative the binding face of XD. In the second encounter complex state N_TAIL_ adopts a more ordered set of conformations with substantially more helical content than is observed in its unbound state and predominantly binds XD in a single orientation that is “upside-down” relative to its orientation in the native complex. This upside-down binding pose is stabilized by the insertion of N_TAIL_ residue L495 into a non-native binding site in the hydrophobic binding groove of XD.

We highlight that while N_TAIL_ conformations in the more disordered N_TAIL_:XD encounter complex state have similar helical propensities to unbound conformations of N_TAIL_ and N_TAIL_ conformations in the more ordered encounter complex state have similar helical propensities to those observed in the native N_TAIL_:XD complex, the deep MSM does not suggest the presence of parallel “induced-fit” and “conformational selection”-type pathways. Transitions from both encounter complex states to the most native-like bound states proceed through similar pathways, illustrating that helical content formed early in folding-upon-binding transitions paths is not necessarily indicative of a conformational selection mechanism. This result is consistent with a previous 1D reaction coordinate transition path analyses of N_TAIL_:XD folding-upon-binding where it was observed that helical content formed early in transition paths frequently breaks to enable the formation of additional native intermolecular contacts before refolding.^56^

There is substantial experimental and computational evidence demonstrating that many IDPs maintain significant conformational disordered when bound to their physiological interaction partners.^23–25, 57^ This phenomenon is frequently referred to as the formation of a “fuzzy” protein complex, and is often explained using the energy-landscape theory inspired concept of conformational frustration.^26, 57, 110–115^ Conformational frustration describes the existence of multiple competing favorable interactions that cannot be simultaneously satisfied and therefore result in a dynamic equilibrium between distinct conformational states. While the existence of fuzzy complexes and the role of conformational frustration in these complexes is well appreciated, few studies have provided atomic resolution molecular mechanisms that rationalize the kinetics of the conformational transitions among the conformational states of IDPs in fuzzy complexes.^55, 57, 67, 98^ The MSMs reported here identify a network of conformationally frustrated bound states of the N_TAIL_:XD complex that share a core set of native intermolecular contacts and are differentiated by the sequential formation of non-native intermolecular and intramolecular contacts that facilitate the folding of additional helical turns. Our analyses provide atomic resolution descriptions of conformationally frustrated states of an IDP in a fuzzy protein complex and quantitative estimates of the time scales of transitions between these states. Our results underscore that an interplay between native intermolecular contacts, non-native intermolecular contacts, and non-native intramolecular contacts produce kinetic barriers between conformationally frustrated states of an IDP in a fuzzy protein complex.^116, 117^ The insights generated from this study and future atomistic studies of fuzzy IDP complexes may ultimately facilitate the design of conformationally frustrated protein complexes with rationally tunable binding affinities.

It was previously noted^56^ that the folding-upon-binding pathways observed in the MD trajectory analyzed here are broadly consistent with previously reported NMR experiments^87^, stopped-flow kinetics measurements^33^ and φ-value analyses of measles virus N_TAIL_:XD binding.^89^ Stopped-flow kinetics measurements clearly resolve separate rates for the formation of an initial encounter complex between N_TAIL_ and XD and the subsequent folding of N_TAIL_^33^, and protein engineering φ-values indicate that encounter complex formation is mediated by hydrophobic residues (A494,L495,L498, and A502) in the central helix of N_TAIL_.^89^ While the simulation analyzed in here was run at higher temperature (400 K) than previous experimental investigations, the MSMs derived in this investigation are broadly consistent with these previously published experimental data.

A recent study of measles virus nucleoprotein and phosphoprotein interactions underlying liquid-like phase separation reported a small set of ^15^N NMR relaxation dispersion data to characterize the binding equilibrium of the measles virus N_TAIL_:XD complex^91^. These data were well fit by a 2-state binding model, suggesting that only one dominant kinetic barrier is resolved in these NMR experiments. As MSMs reported here were derived from MD simulations performed at 400 K and NMR measurements in this experimental investigation were performed at 298K, it is not possible to directly compare the simulated and experimentally measured rates and state populations in these two studies. Building MSMs of N_TAIL_:XD binding at physiological temperatures by combining the VAMPnet approach developed in this work with adaptive sampling strategies could, however, enable a direct comparison between simulated and experimental rates in this system. The recently developed augmented Markov model formalism, where MSM state populations and transition rates are refit using maximum-entropy methods to match agreement with experimental data, provides an eloquent approach to assess the agreement between MSMs and NMR relaxation data.^118^ Such studies may illuminate deficiencies in current molecular mechanics force fields used to study IDP folding-upon-binding, and ultimately facilitate the design of fuzzy protein complexes between IDPs and structured binding partners.

It is interesting to consider the conformational properties of the native-like bound states of the measles virus N_TAIL_:XD complex resolved in this study in the context of previously reported NMR relaxation dispersion measurements used to characterize the binding mechanism of the homologous sendai virus N_TAIL_ molecular recognition element to the homologous sendai phosphoprotein X domain (sendai XD) in unprecedented detail.^22^ In this study, unbound sendai N_TAIL_ was found to be in equilibrium with two bound states, with a population ratio of ∼3:1, that were characterized by chemical shift differences with the unbound state. The more populated bound state was found to contain an elevated population of helical elements relative to apo sendai N_TAIL_ (as assessed by large changes in backbone carbon chemical shifts) but to remain relatively nonspecifically bound (as assessed by relatively small changes in nitrogen and proton backbone chemical shifts in residues at the sendai N_TAIL_:XD binding interface). The less populated bound state has NMR chemical shifts consistent with the fully folded and ordered sendai native N_TAIL_:XD complex. The authors of this study note that the NMR relaxation dispersion data reported are insufficient to provide atomic resolution descriptions of these states, and do not contain information on the relative position of N_TAIL_ on the surface of XD in the more populated bound conformation. This lack of information makes it challenging to understand the microscopic nature of the kinetic barriers between these states.

It is important to caveat that there are substantial differences in the sequences of N_TAIL_ and XD in the sendai and measles viruses. The α-helical molecular recognition element sendai N_TAIL_ has more charged residues (9) than the α-helical molecular recognition element of measles virus N_TAIL_ (5) and the measles virus N_TAIL_:XD binding interface is substantially more hydrophobic than the sendai N_TAIL_:XD binding interface, suggesting that electrostatics and polar interactions are likely to play a larger role in the sendai N_TAIL_:XD binding mechanism.^22, 87^ While one expects there will be appreciable differences in the binding mechanism and bound ensembles of sendai N_TAIL_:XD and measles virus N_TAIL_:XD complexes it is interesting to speculate that the experimentally observed kinetic barriers observed in the bound states of the sendai N_TAIL_:XD complex may share some features with the kinetic barriers identified here. The network of measles virus N_TAIL_:XD bound states reported here contains kinetic barriers that result from the formation of non-native intermolecular and intramolecular contacts that must be broken to facilitate the formation of the fully folded native complex. An analogous set of interactions, perhaps with greater electrostatic contributions resulting from native and non-native salt bridges that confer greater conformational frustration, may underlie the experimentally observed kinetic barriers between bond states of the sendai N_TAIL_:XD complex. Investigating differences in the binding mechanisms of measles virus N_TAIL_ and Sendai N_TAIL_ will be of interest in future investigations. Accurately describing differences in these binding mechanisms will present a stringent test of the quality of MD force fields used to study IDP folding-upon-binding.

Lastly, we have demonstrated the utility of a multi-input neural network framework for describing the conformational dynamics of a highly dynamic intrinsically disordered protein. The approach presented here, where convolutional neural network layers are utilized to reduce the dimensionality of interatomic distance matrices while fully connected dense neural network layers are used to process lower dimensional order parameters describing the helical content of an IDP before combining all features in a fully connected dense neural network, provides a high degree of flexibility for identifying optimal combinations of molecular feature sets with different inherent dimensionalities and embeddings. We demonstrated that this approach distinguishes several structurally and kinetically distinct Markov states that were not resolved using the traditional linear dimensionality reduction tICA approach. We speculate that the deep learning strategy employed here may provide a generalizable approach for learning low dimensional representations of high dimensional IDP simulation data that are best described by multiple distinct degrees of freedom. We plan to investigate the utility of this approach for building MSMs of monomeric IDPs and for identifying collective variables for enhanced sampling methods and diffusion models in future studies.

## Methods

### Markov State Models

Markov state models (MSMs) are stochastic dynamical models that approximate the kinetics of molecules as memoryless, probabilistic jump processes between sets of states.^62^ MSMs utilize a time reversible transition matrix^119^ containing conditional probabilities of transitioning between states. The transition matrix of a MSM is reversible and functions as a transfer operator that propagates a distribution of states, *p*(*t*), forward (and backward) in time by *k*τ discrete steps where k is a positive integer and τ is the lag time of the model.

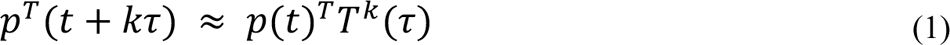

The optimal lag time of a MSM can be determined by ploting the implied time scales (ITS) as a function of the lag time and choosing the lag time at which the implied time scales^120^ are approximatly constant^62^. Additionally, the time resolution of the model can be determined by checking that ITS are above the lag time at which the model is estimated (Supplementary Figures 2 and 11). Implied time scales are determined from the eigen values, λ_i_, of the transition matrix.

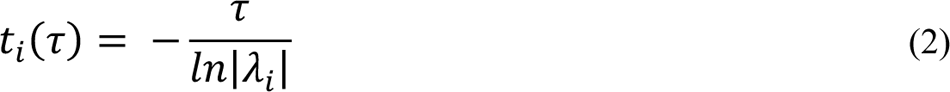

By definition, MSM transition matrices have a maximum eigen value of 1 whose eigen vector corresponds to the steady state or stationary population, π, of states as time approaches infinity.^121^

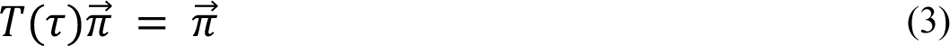

Using the the stationary distribution and the transition matrix of a MSM, the mean first passage times between pairs of states (*MFPT_ij_*) can be determined from an N_states_ by N_states_ system of equations (Supplemenary Figures S9 and S17).^122^

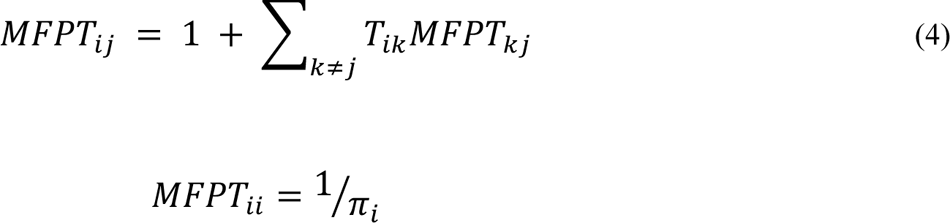

In addition to ITS, validation of MSMs and their transtion matrices is determined by the Chapman-Kolmogrov equation^62, 121^,

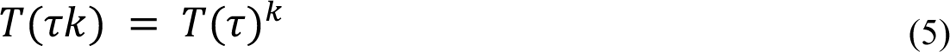

in which the ability of a transition matrix to reproduce transistion probabilities at longer timesales is evaluated (Supplementary Figures 2 and 11).

### Input Data for Markov State Models

We utilized the 200 μs unbiased MD trajectory from Robustelli et. al^56^ which contains N_TAIL_ residues 484-504, XD residues 458-506 and 20mM of NaCl in a 72 Å per side cubic box. This trajectory was parametrized using the a99SB-disp force field, a99SB-disp water model^52^ and contained 1,000,000 frames with a spacing of 200ps. For the construction of our MSMs^62, 70^, we only considered a continuous 167μs subset (from 3μs to 170μs) of the original trajectory in which XD predominantly remains in its folded state. We generated the molecular features for MSM construction and neural network training by calculating intermolecular distances between all residues of N_TAIL_ and XD using the minimum distance between heavy atoms. Additionally, we computed the α-helical order parameter Sα ^102^ and identified helical conformations using the DSSP^101^ algorithm. The order parameter Sα quantifies the helical content of each 7-residue segment of a peptide chain and is computed by the following,

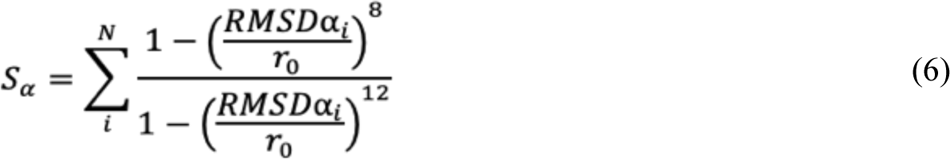

where RMSDα_i_ denotes the root mean squared deviation between each 7-residue segment of N_TAIL_ and a geometrically perfect alpha helix comprised of the same residues. The exponential terms in the equation act as a switching function to output values between 0 (not helical) and 1 (perfectly helical) for each segment. The threshold of the switching function is tuned by the parameter r_0_, which was chosen to be 0.8 Å. Setting the parameter r_0_ to 0.8 Å has the effect of reducing RMSDα_i_ values > 2.5 Å to nearly zero and RMSDα_i_ values < 0.5 Å to nearly 1. For constructing MSMs, we chose to omit the summation in (eq. 6) to retain a more localized description of the helical content of N_TAIL_. As a result of the 7-residue sliding window used in the computation of Sα and N_TAIL_ being 21 residues long, we compute a length 15 vector for each time step of the simulation describing the helical content of every possible contiguous 7 residue segment of N_TAIL_. We note that for broad statistical characterizations (such as in Figure 1), the summation in equation 1 is retained to provide an estimate of the total helicity of N_TAIL_ per simulation frame (“N_TAIL_ Sα”).

We constructed the second α-helical descriptor for N_TAIL_ using the DSSP Algorithm. The DSSP algorithm uses dihedral angles and hydrogen bonding analysis to classify the secondary structure of each residue in a peptide chain. The secondary structure predictions given by DSSP were then numericized by equating helical classifications to 1 and all others to zero. As a result, the processed binary DSSP assignments produce a vector of length 21 for each time step of the simulation with values indicating if each residue of N_TAIL_ is in a helical conformation (value of 1) or not (value of 0). Both Sα and binary DSSP features were considered in quantifying the helical content of N_TAIL_ as they evaluate helical content using distinct metrics and as a result, produce differing degrees of locality in the descriptions they provide.

We tested several feature sets for state discretization including combinations of interatomic distances, dihedral angles, fraction native intermolecular contacts (*Q*), binary DSSP assignments and Sα values. We assessed the quality of feature sets by comparing VAMP2 scores^76, 83^, the spectral gap observed among the eigenvalues of the dominant tICA eigenmodes,^79, 96, 97^ and the ability of each feature set to resolve conformationally distinct free energy basins in low dimensional tICA projections. For tICA, we found the combination of intermolecular residue distances and Sα best satisfied these metrics and that the addition of DSSP features had negligible effect. We subsequently omitted the DSSP features from our tICA analysis and used only intermolecular distances and Sα order parameters. In contrast, we found that including DSSP features in our VAMPnet increased the model’s ability to differentiate N_TAIL_ conformations differing only in the helical content of residues near the termini; thus, we used a feature set containing intermolecular residue-residue distances, Sα, and binary DSSP helical assignments as input data in our VAMPnet implementation.

### Construction of a hidden Markov state model (HMSM)

To construct an initial MSM, we performed tICA on a feature set comprised of the nearest-heavy-atom intermolecular distances between all residues of N_TAIL_ and XD and Sα values. The tICA lag time, number of tICA components (tICs) used for clustering, and the number of *k-means* clusters were optimized based on the interpretability and distinctness of the structural properties of the resulting clusters. We iteratively computed tICA with varying lag times and clustered the resulting tICs using a varying number of components and k-means clusters. We characterized the structural properties of clusters by computing their distributions of the fraction of native intermolecular contacts (*Q*), N_TAIL_ *Sα*, Radius of gyration (R_g_), intermolecular contact probabilities and helical assignments from the DSSP algorithm. We found that using a lag time of 6 ns for tICA, clustering conformations using the ten time independent components (tICs) with the largest eigenvalues and implementing *the k-means* algorithm with seven cluster centers produced the most interpretable and conformationally distinct clusters. However, upon estimating MSMs from these clusters over a range of lag times, we found that for lag times up to 24 ns, these models produced resolved, but non-converged implied timescales (data not shown). These MSMs also failed to reproduce transition probabilities for non-native bound states at longer timescales.

To produce MSMs with both converged time scales and robust CK-tests, we employed hidden Markov state models (HMSMs). HMSMs are an effective tool for building robust and reproducible MSMs for high dimensional systems where finding a set of Markov states that pass validation tests is challenging.^95^ Projected HMSMs are estimated from transitional MSMs; the slowest relaxing timescales of the original MSM are used to coarse grain its states to a smaller number of metastable sets. The number of metastable sets used to build an HMSM should be equal to or less then the number of resolved timescales in the conventional MSM they’re estimated from. We built our HMSM by estimating a series of HMSMs from MSMs with varying numbers of states and lag times. We increased the number of states in the initial MSMs by employing the k-means clustering algorithm with larger numbers of centroids to cluster the same ten tICs we previously found to be optimal to prevent the HMSM coarse graining from reducing our model to too few states. We found that using a lag time of 6 ns, twelve initial clusters and coarsening to seven states produced robust HMSMs (in terms of timescales and CK-tests) with the fewest number of states (Supplementary Figure 2).

### Unconstrained VAMPnet and neural network architectures

The feature set used to train the deep MSM was comprised of the intermolecular distances between all residue pairs of N_TAIL_ and XD, Sα order parameters and binary DSSP assignments. We employed a multi-input deep learning approach where each feature type was processed separately before being aggregated with the other features to make state predictions. This approach allows for the input feature set to be optimized internally and each feature type to be processed using neural network layers that best suit its inherent data structure. This approach enabled us to treat the matrix of intermolecular distances (or “contact map”) calculated in each frame of the simulation as an image and utilize convolutional neural network layers to leverage the local spatial coherence in this representation. We utilized separate sets of fully connected neural network layers to process the Sα and binary DSSP feature sets. Each instantaneous set of intermolecular residue distances were arranged into a 49 by 21 matrix where each index represents the intermolecular distance between each residue in XD (49 residues) and N_TAIL_ (21 residues). Each set of Sα and binary DSSP values were placed into length 15 and 21 vectors, respectively. In aggregate, the VAMPnet dataset is comprised of 3 distinct feature sets, each processed separately by distinct sets of neural network layers (or lobes), before being aggregated and transformed through a final lobe, containing fully connected neural network layers (Figure 2). The output of the final lobe is capped with a SoftMax activation function to produce a normalized distribution that describes the probability of a frame being assigned to each Markov state.

We determined the architecture of our neural network by varying the number of layers and their widths in each lobe of the neural network. To reduce computational overhead, we constrained our optimization of the neural network architecture by requiring that each lobe contain the same number of layers and that the lobes used to transform the N_TAIL_ Sα and DSSP helical order parameters be identical apart from their input layers. In addition, the possible configurations of the convolutional layers used to transform intermolecular distance matrices were constrained based on the shape the input (49 XD residues by 21 N_TAIL_ residues). We determined our architecture by first performing a grid search over a range of configurations and then performed a Bayesian optimization around the optimal parameters identified in the initial grid search. For the Bayesian optimization, we used the tree-structured Parzen estimator algorithm^123, 124^ implemented in the *optuna*^125^ software. A detailed diagram of the final neural network architecture determined from the Bayesian optimization procedure is displayed in Supplementary Figure 10. After determining the neural network architecture, we employed this procedure to determine the optimal batch size, optimizer learning rate and epsilon parameter. We found that using learning rate of 5e-6, a batch size of 16384 and an epsilon parameter of 1e-7 produced optimal results.

Additional hyperparameters of VAMPnets include the lag time of the model and the number of output states. To determine these hyperparameters, we conducted optimization runs incrementally increasing the values of each hyperparameter while holding the other hyperparameters constant. We judged the success of these trials based on the maximization of the VAMP score relative to its highest possible value and the interpretability of the learned state assignments in terms of the fraction of native contacts (*Q*), Sα, radius of gyration and RMSD from the native complex. We found that using 12 output states and a lag time of 2 ns to train the unconstrained VAMPnet best satisfied these conditions and consistently produced similar sets of states. The final architecture of multi-input neural network used in our VAMPnet implementation is shown in Supplementary Figure 10.

We trained our initial unconstrained VAMPnet using the VAMP2 score. The VAMP2 score evaluates the so-called kinetic variance between each neural network transformed sample, *X*_0_(*X*_t_), of the dataset and it’s time-lagged analogue, *X*_τ_(*X*_t+τ_), where *X*_0_ and *X*_τ_ are neural network transformations that convert molecular features into probabilistic Markov state assignments and *X*_t_ and *X*_t+τ_ are instantaneous sets of molecular features at times t and t+τ.^84^ Optimizing the VAMP2 score of transformations *X*_0_(*X*_t_) and *X*_τ_(*X*_t+τ_) is analogous to solving the problem of finding orthonormal transformations of *X*_t_and *X*_t+τ_with maximal time-correlations and corresponds to finding the best linear approximation^84^ to the following,^83^

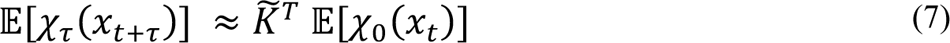

where *K*^T^ is the finitely estimated Koopman matrix that transforms a potentially non-linear dynamical system or dataset into a latent space which, on average, evolves linearly in time. The VAMP2 score is defined as the Frobenius norm or sum of the squared singular values (σ_i_) of the half-weighted Koopman matrix, *C_00_^−½^C_0τ_C_ττ_^−½^*

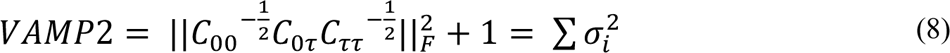

Where the covariance matrices, C_00_, C_0τ_ and C_ττ_ are defined by mean free neural network transformed instantaneous and time lagged data as follows.

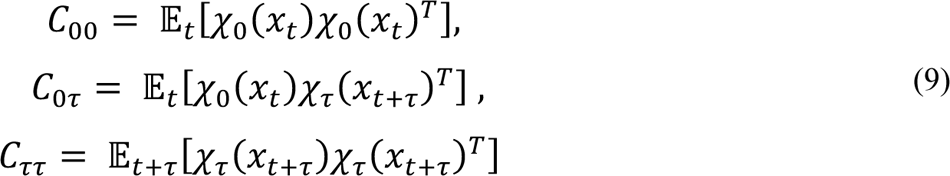

We note that in general, neural network transformations, *X*_0_ and *X*_τ_ can be distinct neural network architectures with independently trained weights, however, in our implementation *X*_0_ ≡ *X*_τ_.

### Training the constrained VAMPnet to construct a deep MSM

After determining the optimal architecture and hyperparameters for the unconstrained VAMPnet, we proceeded to build a constrained VAMPnet using the same architecture with the addition of two constraint layers. In the constrained VAMPnet^85^, the constraint layers (*u* and S) are implemented to ensure the learned transition matrix is both stochastic (all positive elements) and reversible (obeys detailed balance). Constraint *u* is a vector of length equal to the number of states used to weight data towards equilibrium and constraint S is matrix of shape N_states_ by N_states_ used to estimate a reversible transition matrix. The constrained VAMPnet was trained with a modified version of VAMP-E score that incorporates the constraints *u* and S.

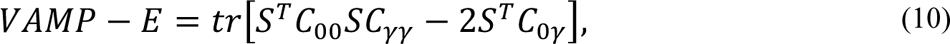

where

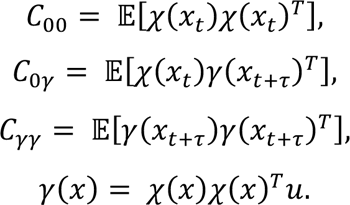

Here, gamma is a weighted state representation used to compensate for non-equilibrium state assignment probabilities. We trained our constrained VAMPnet 30 separate times starting from the same initial unconstrained VAMPnet.

In the constrained VAMPnet procedure, both the weights of the unconstrained VAMPnet and constraint layers are optimized, thus, retraining only the constrained VAMPnet also modifies the weights of the initial, unconstrained VAMPnet. We note that using the same unconstrained VAMPnet in each optimization of the constrained VAMPnet produces small error estimates that may be underestimated compared error estimates obtained from retraining the unconstrained VAMPnet multiple times. Given the large number of parameters in our neural network architecture (∼4e6 parameters), we used this approach to circumvent considerable computational costs and consider these error estimates as lower bounds of the trial errors. As outlined in its original implementation^85^, it is recommended to include an initial step in which only the constraints of the constrained VAMPnet are trained using batches containing all training data. When training the unconstrained VAMPnet and the constraints together (a separate step), we attempted to stay consistent with this strategy and used the largest batch size possible given our computational resources which was 56,000 time-lagged pairs of data. To estimate the implied timescales and CK-tests, we retrained only the constraints of the constrained VAMPnet at integer multiples of the initial lag time (6 ns) which was done for all 30 optimization runs. We chose to use a lag time of 6 ns for the constrained VAMPnet based on the results of these validation measures which we found to produce the most reproducible and robust models in a series of initial estimations of the constrained VAMPnet at varying lag times (Supplementary Figure 11).

### Neural network training

In both the unconstrained and constrained VAMPnets, we used a 9:1 train-validation split, randomly shuffled time lagged pairs of data and implemented early stopping to prevent overfitting where we saved network weights each time the VAMP score reach a new maximum. We implemented all neural networks in using the deep learning library *PyTorch*^126^.

### Estimation of trajectory observables and error analysis

For the HMSM, all MSM observables and error estimates were computed using the *pyemma*^70^ and *deeptime*^127^ software packages via Bayesian hidden markov models which use a gibbs sampling scheme to resample the transition matrix. Here, we estimated errors by resampling the HMSM transition matrix using 100 trials. All HMSM trajectory observables are the bootstrap mean and its associated 95% confidence intervals computed from the results of the resampling procedure. For the deep MSM, we trained the final model using 30 independent trials and computed both MSM and trajectory observables from the trained models. All statistical analysis of the trajectory observables of the deep MSM states and MSM observables are computed by bootstrapping / aggregating the results of these 30 trials, e.g. average values, 95% confidence intervals of averages, standard deviations, weighted historgrams and discrete probability distributions. Trajectory observables from the deep MSM states were computed from the probabilistic state assignments produced from each optimization run by the following,

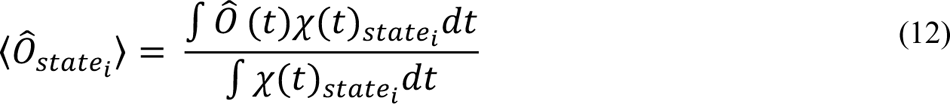

where Ô(t) represents an arbitrary trajectory observable computed for every frame (t) of the trajectory and *χ*(*t*)_statei_ is a probabalistic state assignment for every frame (t) of the trajectory. Using this definition, we can also compute the standard deviation of trajectory observables by the following equation.

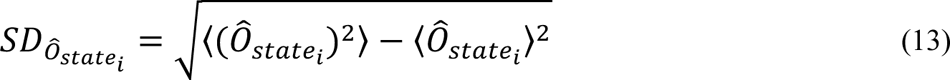

We combine uncertaines computed from separate trials and contact popualtions for different residue pairs by combining variances,

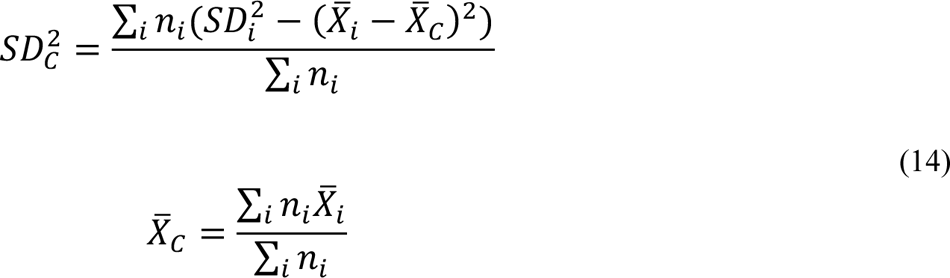

Where *SD*^2^_c_ is the combined variance, *n*_i_ are the number of trials used to compute the mean and standard deviation of each statistic to be combined, *X*_i_ are the means of each statistic to be combined and *X*_c_ is the combined mean.

### Fraction of Native Intermolecular Contacts

The fraction of native intermolecular contacts (*Q*), as defined in Robustelli et al^56, 100^, was used to characterize the formation of the N_TAIL_:XD complex. The fraction of native contacts at each simulation time step, (t), was calculated by the following,

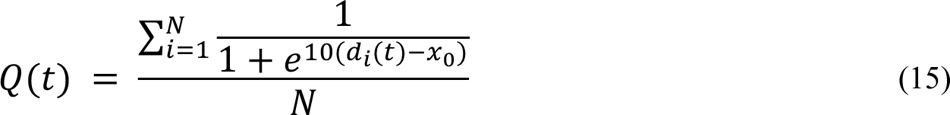

where d_i_ represents the nearest neighborh heavy atom distance between each pair of native contacts, N is the total number of native contact pairs and *X*_&_ is a cutoff distance of 5Å. Native intermolecular contacts were previosuly defined as those contacts which remained stable (populated > 80%) in an MD simulation of the native N_TAIL_:XD complex run at 400 K, to match the temperature of the equilibrium folding-upon-binding simulation analyzed here.^56^

### Color gradients of structural snapshots

We computed the color gradients of the structural snapshots of N_TAIL_:XD using a modified version of the fraction of native intermolecular contacts based only on the crystal structure of the native complex (PDB 1T6O).^86^ For establishing color gradients, we defined native contacts as any intermolecular residue pair between N_TAIL_ and XD with a minimum heavy atom distances less than 5 Å in PDB 1T6O. Correspondingly, we define non-native contacts for each residue as all other possible intermolecular contacts that have not been identified as native. In each simulation frame two residues are considered to be in contact if their nearest heavy atom distance is less than 5 Å. We compute the average population of the native and non-native contacts of every residue in each Markov state. For coloring structures, we normalize native and non-native fractions by dividing each by the largest fraction observed in any Markov state (∼ 0.99 and ∼0.14 for native and non-native fractions, respectivly) which assigns a value between 0 and 1 for each residue in each Markov state. We then set a color gradient ranging from 0 to 1 in the molecular visualization software *pymol*^128^and set the beta value of each residue (alpha carbon) to the normalized fraction of native and non-native contacts. The normalization step allows the scale of the color gradients to be the same across all structures, thus allowing for quantitative comparision of the contact profiles of each Markov state via their structural snapshots.

## Data & Code Availability

All code used for trajectory analyses and the construction and validation of the hidden Markov state model and deep Markov state model are freely available from GitHub (https://github.com/paulrobustelli/Sisk_NTAIL_DeepMSM_2023). The 200 μs N_TAIL_:XD MD trajectory analyzed here is available for non-commericial use by request from D.E. Shaw Research.

## Acknowledgements

This work was supported by the National Institutes of Health under award R35GM142750.

