## Supplemental Information for "Folding-upon-binding pathways of an intrinsically disordered protein from a deep Markov state model"

† To whom correspondence should be addressed.

Paul Robustelli

Address: 6128 Burke Laboratory

Department of Chemistry

Hanover, NH, 03755

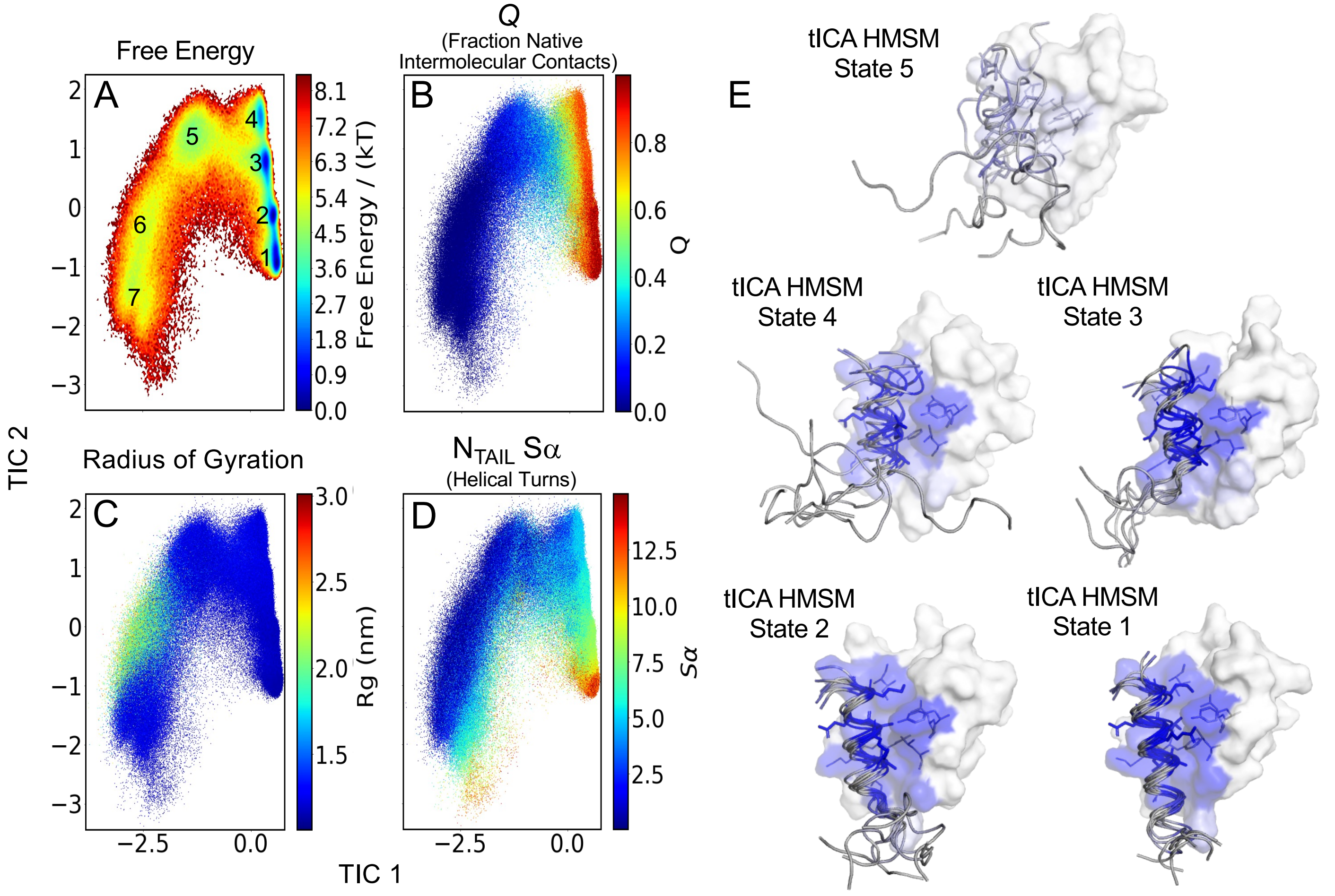


**Supplementary Figure 1. Time-lagged independent component analysis of a long-time scale molecular dynamics simulation of the measles virus nucleoprotein $N_{TAIL}$ reversibly binding the X domain (XD) of the measles virus phosphoprotein complex. (A)** The free energy surface of $N_{TAIL}$:XD conformational states observed in an MD simulation as a function of the two dominant time lagged independent components (TICs) obtained from time lagged independent component analysis (tICA). The molecular features computed from each frame of the $N_{TAIL}$:XD MD simulation are projected onto the tICA surface, and each projected point is colored by the fraction of native intermolecular contacts $Q$ **(B)**, the radius of gyration of all Cα atoms of $N_{TAIL}$ and XD **(C)** and the value of $N_{TAIL}$ Sα **(D). (E)** Structural representations of native-like bound states (tICA HMSM states 1-4) and the encounter complex state (tICA HMSM state 5) resolved by tICA followed by hidden Markov state model (HMSM) state partitioning. Each tICA HMSM state representation is an overlay of multiple representative $N_{TAIL}$ structures with one surface representation of XD. The residues of $N_{TAIL}$ and XD are colored by a gray-to-blue gradient that represents the state-averaged fraction of native intermolecular contacts for each residue. Darker blue coloring indicates residues with the higher state-averaged fractions of native intermolecular contacts.

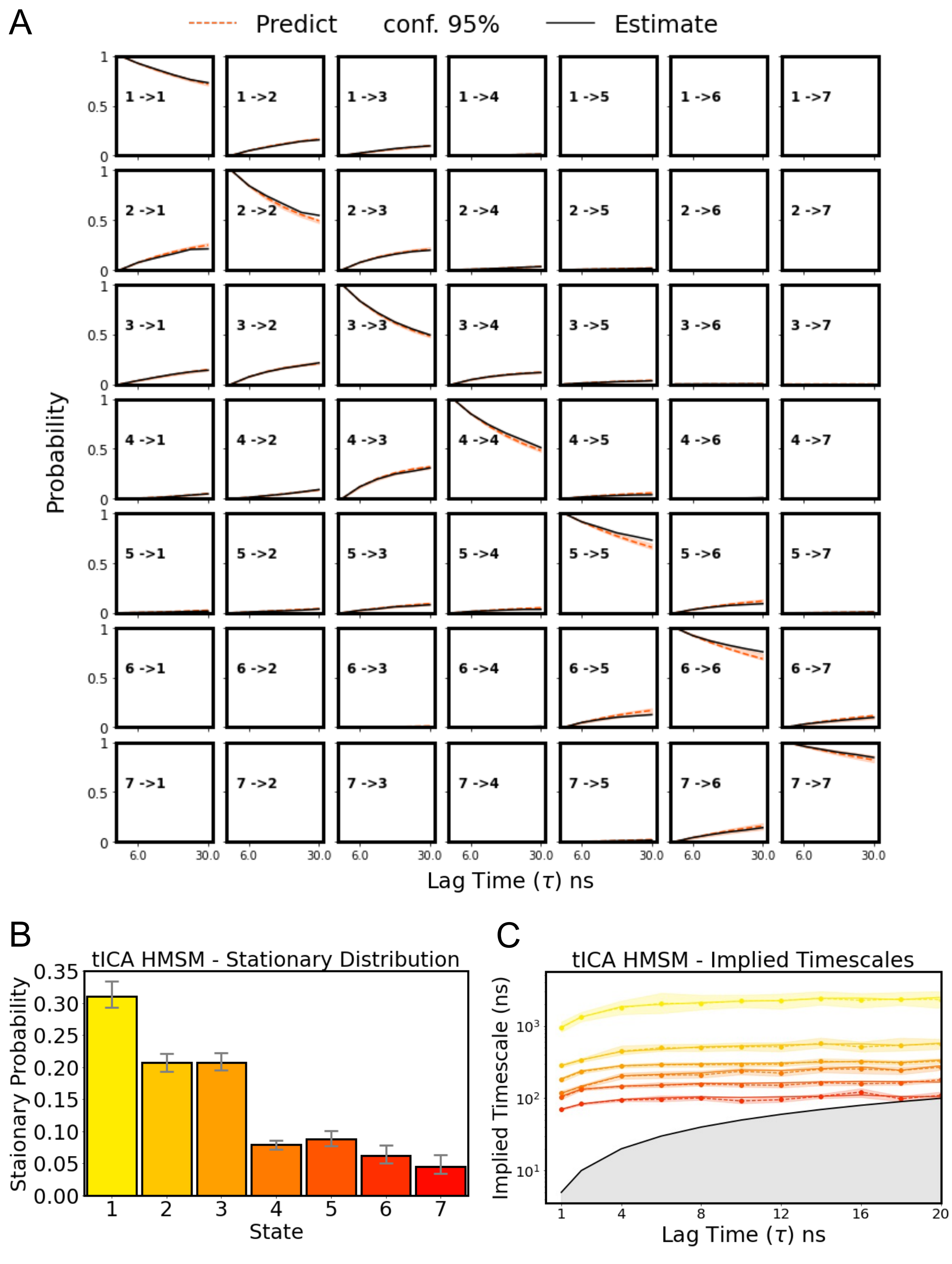


**Supplementary Figure 2. tICA HMSM validation tests and stationary distribution. (A)** The Chapman-Kolmogrov (CK) test calculated for the tICA HMSM. The CK test evaluates the dependence of an MSMs predictions on the chosen lag time. The CK-test compares the evolution of transition probabilities for each state, i, to every other state, j, for integer multiples of the initial lag time of the model (6 ns). Orange dotted lines ("Predict") represent transition probabilities predicted by propagating the tICA HMSM transition matrix and the solid black lines ("Estimate") indicate transition probabilities obtained from transition matrices resampled from the trajectory data at integer multiples of the tICA HMSM lag time. The orange shaded region indicates the 95% confidence interval of the mean obtained from Gibbs sampling with 100 samples. **(B)** The stationary distribution for each state of the tICA HMSM with error bars showing the deviation of the 95% confidence interval of the mean obtained from Gibbs sampling of the transition matrix using 100 samples. **(C)** The log scaled implied timescales obtained from the tICA HMSM transition matrices estimated at increasing lag times. The colored shaded regions show the deviation of the 95% confidence interval of the mean of the implied timescales estimated for each lag time obtained from Gibbs sampling using 100 samples. The solid black line and gray shaded region indicate time scales equal to or less than the lag time and represents the threshold for the fastest ITS that can be resolved by the model.

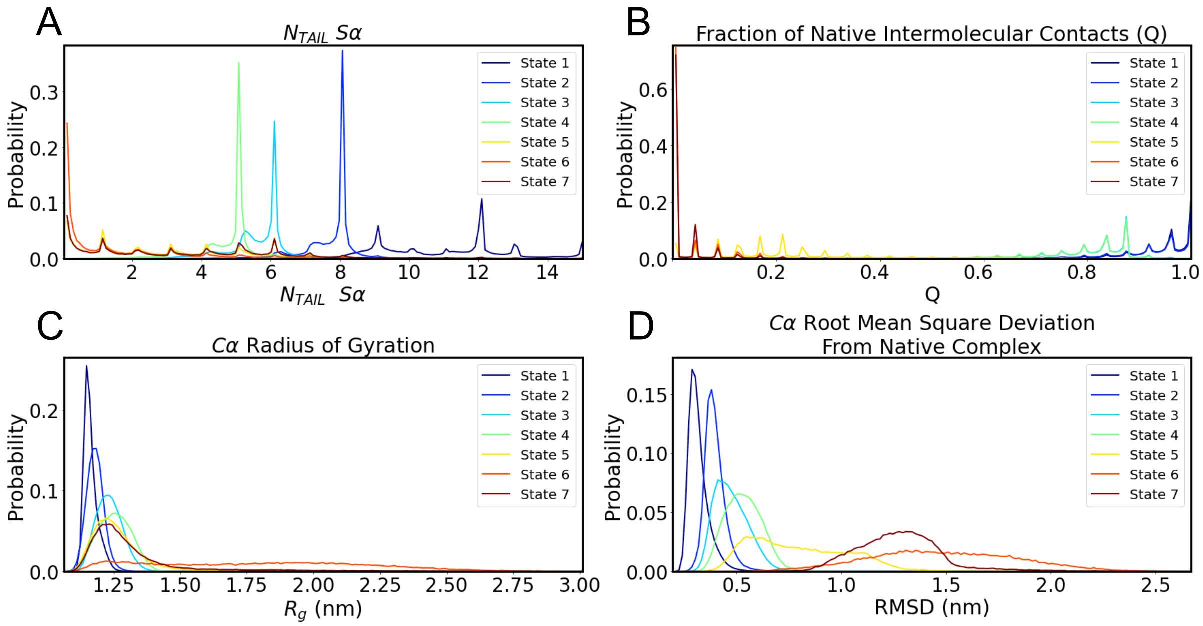


**Supplementary Figure 3. Structural properties of the tICA HMSM states.** The discrete probability distributions of **(A)** $N_{TAIL}$ S$\alpha$ (the number of ideal helical turns in $N_{TAIL}$), **(B)** the fraction of native intermolecular contacts *Q*, **(C)** the radius of gyration computed using all C$\alpha$ atoms of $N_{TAIL}$ and XD and **(D)** the root mean squared deviation of $N_{TAIL}$ and XD C$\alpha$ atoms from a solved X-ray crystal structure of the native complex (PDB 1T6O). The distribution of each statistic is computed separately for each state of the tICA HMSM.

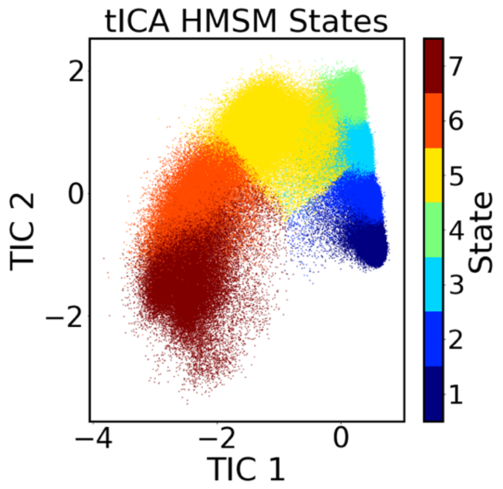


**Supplementary Figure 4. tICA HMSM state map.** HMSM state assignments of each simulation frame projected on to the two dominant time-lagged independent components (TICS) obtained from tICA.

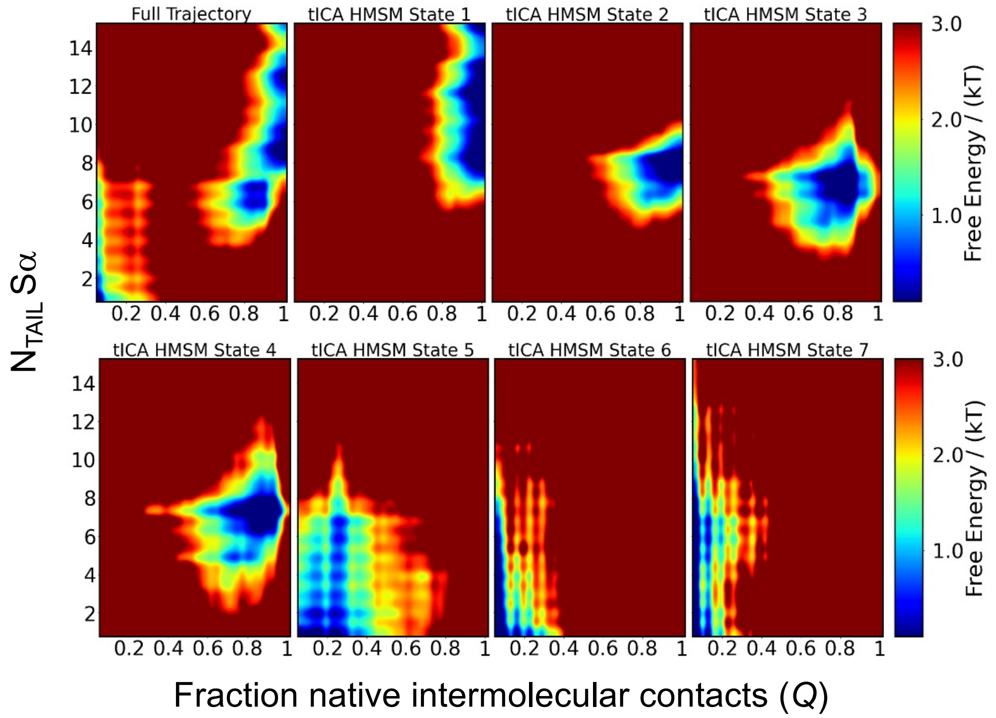


**Supplementary Figure 5. Free energy surfaces of the tICA HMSM conformational states as a function of $N_{TAIL}$ Sα and the fraction native intermolecular contacts (*Q*).** Free energy surfaces of the full MD trajectory and each of the tICA HMSM states shown as a function of the fraction of native intermolecular contacts (*Q*) and $N_{TAIL}$ Sα (the number of ideal helical turns in $N_{TAIL}$).

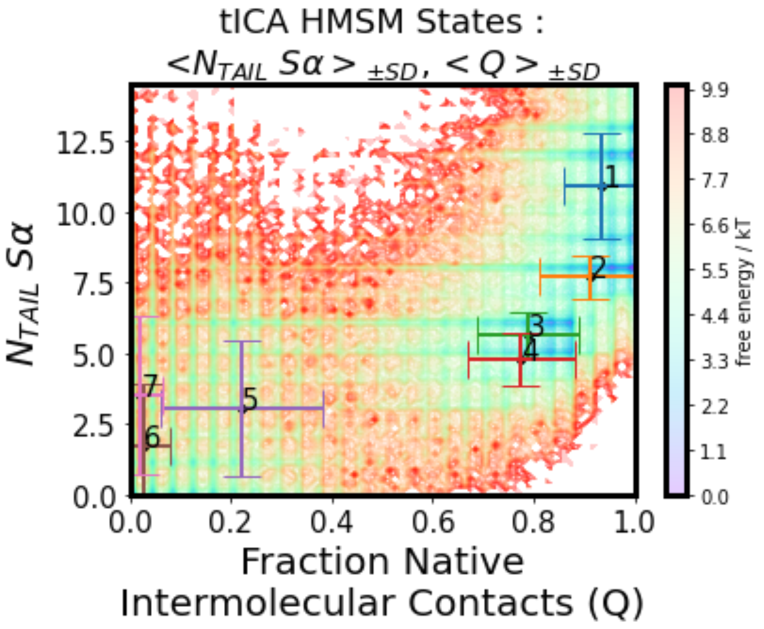


**Supplementary Figure 6. Comparison of properties of tICA HMSM states.** Free energy surface of $N_{TAIL}$:XD complex formation as a function of the α-helical folding order parameter Sα and the fraction of native intermolecular contacts *Q*. The mean values of $N_{TAIL}$ Sα (<$N_{TAIL}$ Sα>) and *Q* (<*Q*>) are indicated for each tICA HMSM state. Error bars represent the standard deviation (SD) of each quantity.

| State | $\langle Q \rangle$ | $Q_{SD}$ | $\langle N_{TAIL}\ S\alpha \rangle$ | $\langle N_{TAIL}\ S\alpha \rangle_{SD}$ | $\langle R_g\ (nm) \rangle$ | $\langle R_g\ (nm) \rangle_{SD}$ |
|---|---|---|---|---|---|---|
| 1 | 0.935 | 0.074 | 10.926 | 1.895 | 1.157 | 0.025 |
| 2 | 0.911 | 0.101 | 7.715 | 0.763 | 1.174 | 0.032 |
| 3 | 0.789 | 0.099 | 5.373 | 1.084 | 1.227 | 0.049 |
| 4 | 0.776 | 0.108 | 4.770 | 0.926 | 1.248 | 0.061 |
| 5 | 0.222 | 0.160 | 3.053 | 2.380 | 1.269 | 0.143 |
| 6 | 0.025 | 0.054 | 1.742 | 2.194 | 1.779 | 0.397 |
| 7 | 0.021 | 0.045 | 3.519 | 2.798 | 1.323 | 0.228 |

**Supplementary Table 1**. **Structural Properties of tICA HMSM States**. The values of the mean and standard deviation (SD) of $N_{TAIL}$ Sα and *Q* for each state of the tICA HMSM.

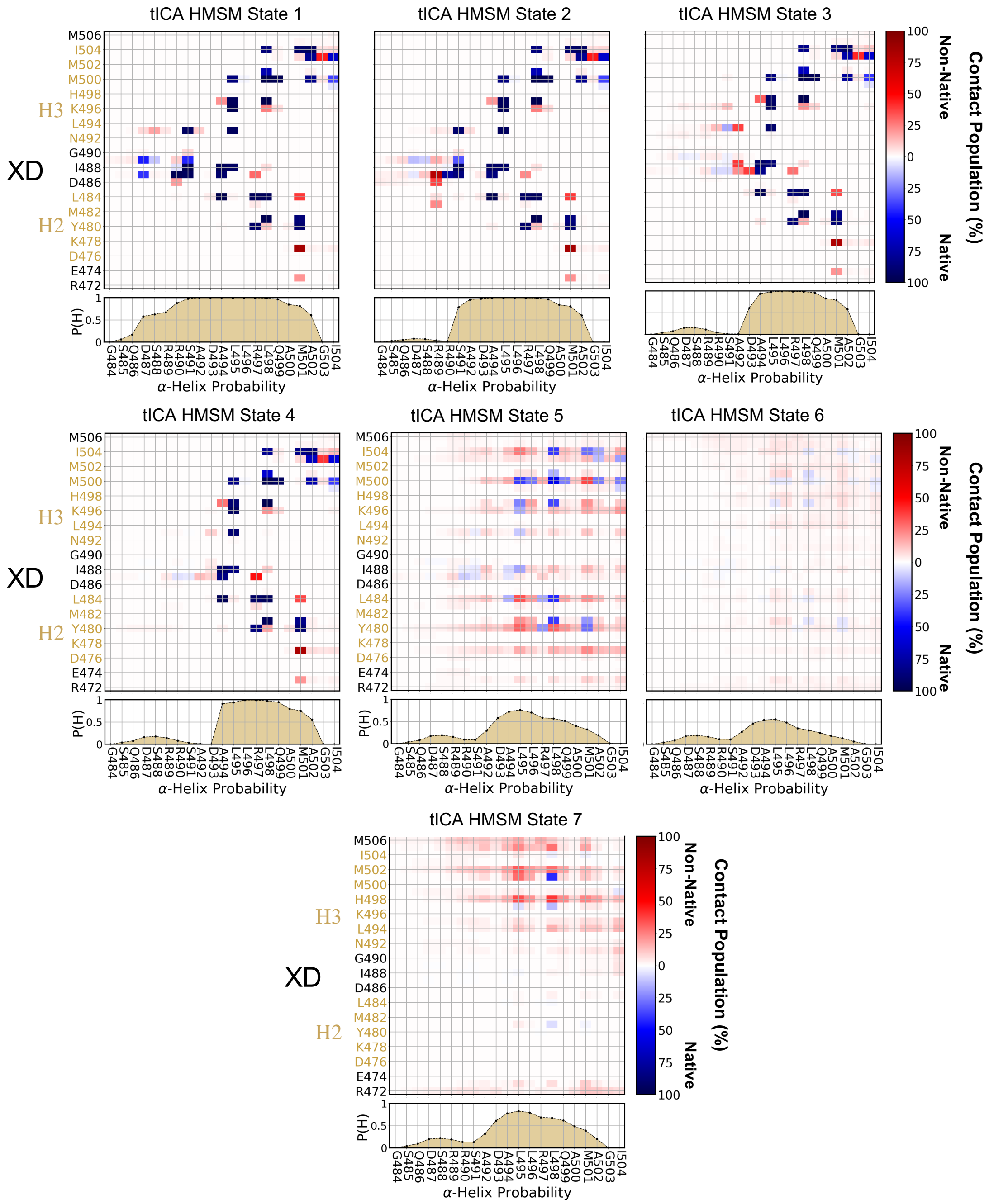


**Supplementary Figure 7. Intermolecular contact populations and $N_{TAIL}$ helical propensities of tICA HMSM states.** State averaged intermolecular $N_{TAIL}$:XD contact populations and $N_{TAIL}$ helical propensities for each tICA HMSM state. Intermolecular contacts were defined as occurring in all frames where the minimum distance between heavy atoms of two residues was less than 5.0 Å. Native intermolecular contact pairs are colored blue and non-native intermolecular contact pairs are colored red**.** Native contacts are defined as those present in the crystal structure (PDB 1T6O) using the same criteria. Helical propensities (P(H)) were calculated using DSSP.

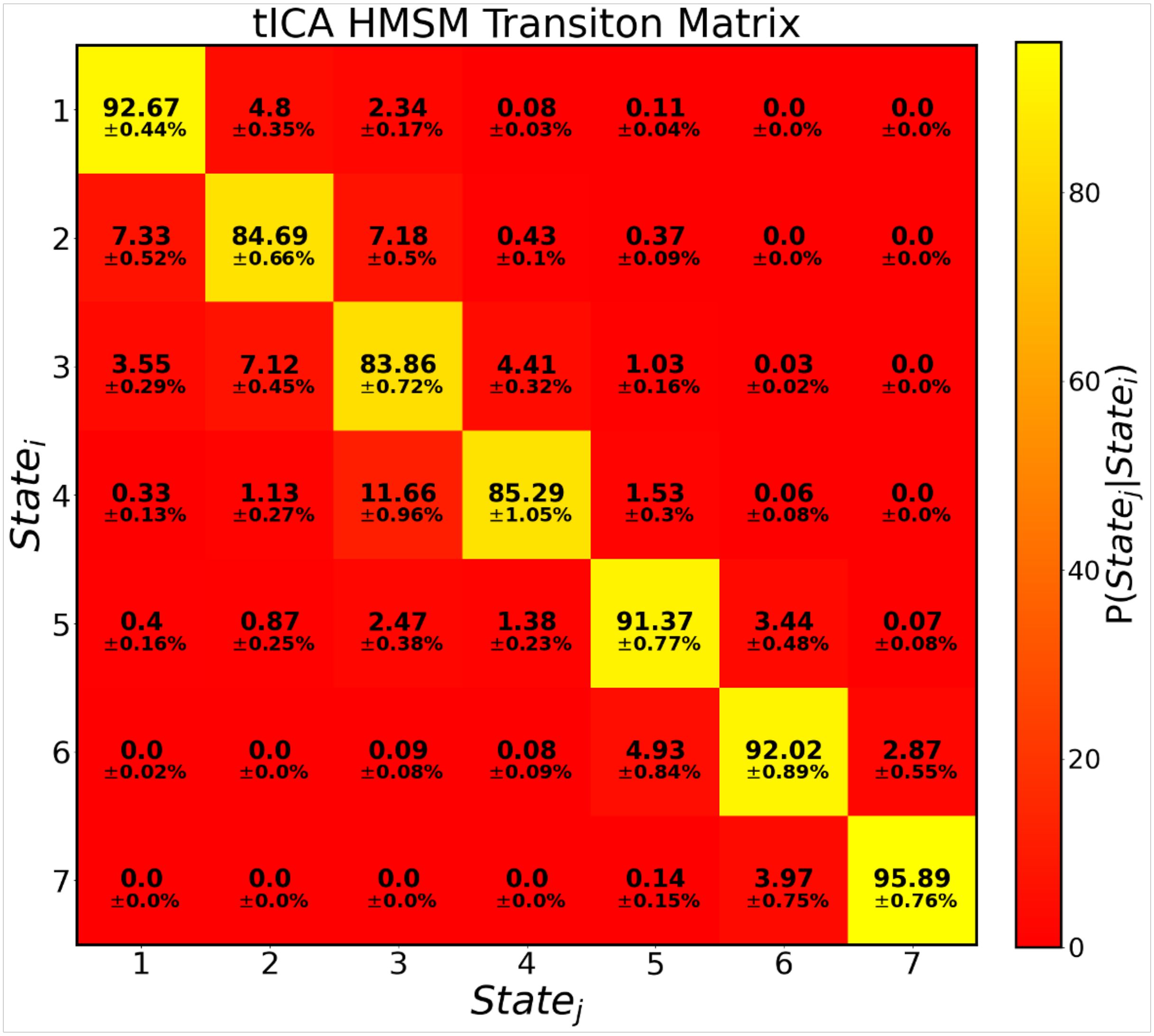


**Supplementary Figure 8. tICA HMSM transition matrix for lag time τ = 6 ns.** The transition matrix for the tICA HMSM estimated at lag time, τ = 6 ns. The transition matrix defines the conditional probability of transitioning from $state_i$ (at time t) to $state_j$ (at time t+ τ). The transition matrix is estimated by counting transitions between states at the given lag time over the simulation trajectory, applying a maximum likelihood estimator to enforce reversibility and then estimating a HMSM. The transition matrix shown here is the bootstrap mean of results obtained from Gibbs sampling of the transition matrix using 100 samples. Errors report the mean of the upper and lower deviations of the 95% confidence interval of the bootstrap mean.

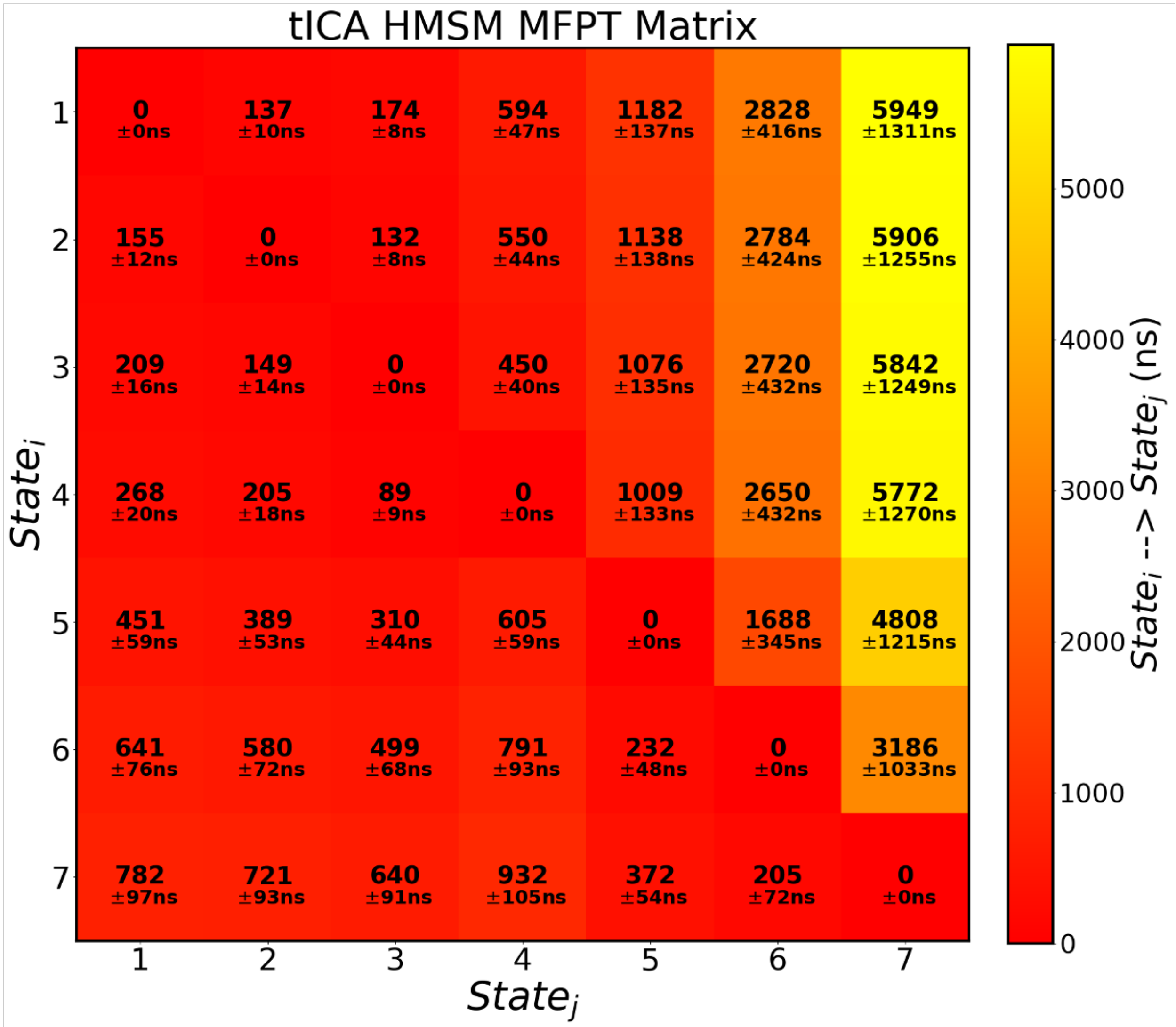


**Supplementary Figure 9. tICA MSM mean first passage time matrix.** The mean first passage time (MFPT) matrix shows the mean time duration it takes to transition between Markov states of the tICA HMSM. We compute the mean first passage times using the transition matrix and stationary probabilities of the tICA MSM estimated at lag time, $\tau = 6$ ns. Errors report the mean of the upper and lower deviations of the 95% confidence interval calculated from Gibbs sampling of the transition matrix using 100 samples.

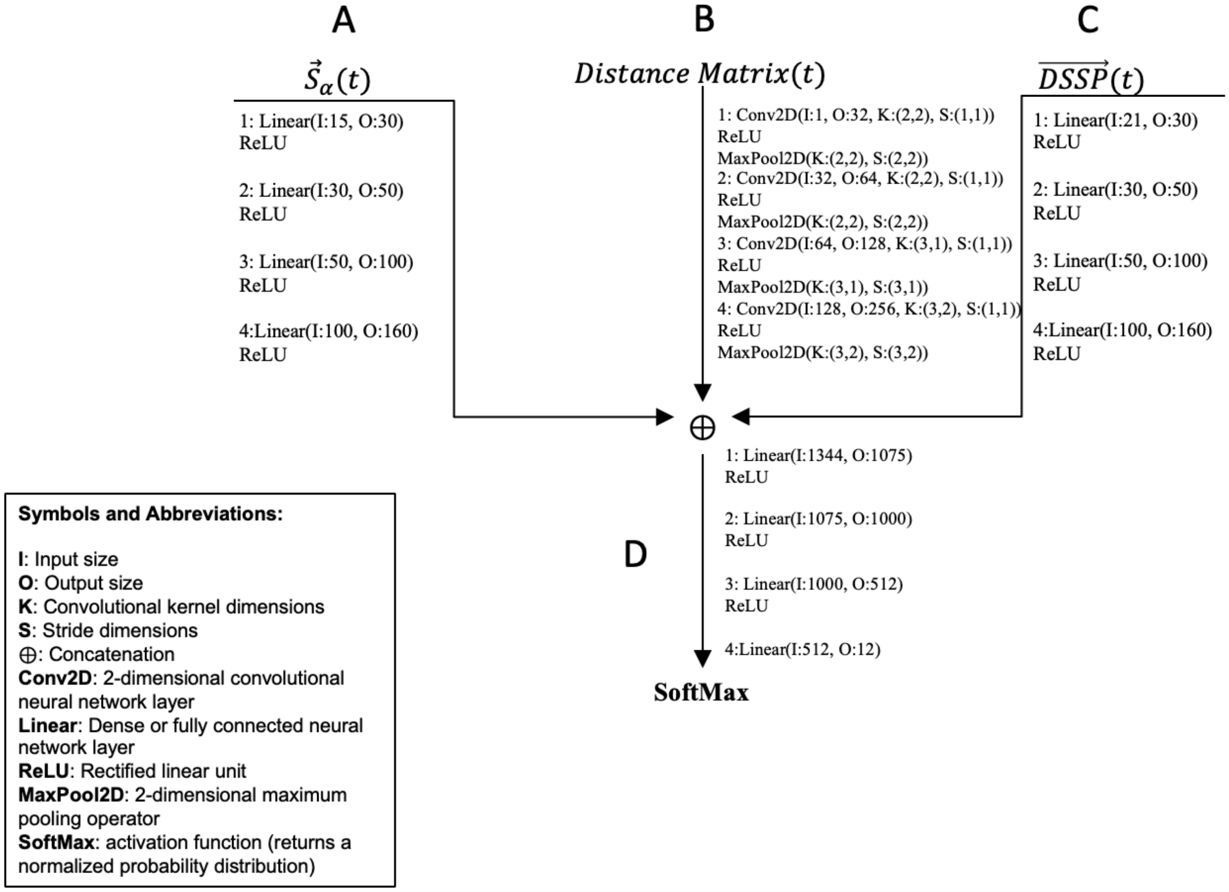


**Supplementary Figure 10. Multi-input Neural Network Architecture.** Schematic diagram of the neural network architecture used in our VAMPnet implementation with all transformations and their corresponding dimensions. **(A, C)** The sub-neural networks used to process the vectorized alpha helical order parameters S$\alpha$ (A) and a binary, helix-only version of DSSP (C). These sub-networks are fully connected (or "dense") neural networks that use rectified linear unit (ReLU) activation functions. These transformations expand the dimension of the helical order parameters to ensure that the network does not focus entirely on the distance matrix due to its considerably higher dimension (there are 1029 intermolecular residue distances in the distance matrix while the S$\alpha$ and DSSP vectors are only length 15 and 21, respectively). **(B)** The convolutional sub-neural network used to process the intermolecular residue distance matrix input. The convolutional neural network layers use progressively larger kernel sizes (rectangular kernels) with respect to the zeroth axis to account for the non-square distance matrix. The distance matrix input is non-square because it contains all intermolecular residue distances between each residue in the XD (49 residues) and $N_{TAIL}$ (21 residues). Following each convolutional neural network layer rectified linear units are employed to introduce non-linearities and max pooling layers, which take the maximum of the transformed distance matrix values in each kernel, are used to reduce the dimension of the distance matrix to lessen the discrepancy between the number of distance and helical features. **(D)** The final sub-neural network used to combine and transform the outputs from the first 3 sub-neural networks is comprised of 4 fully connected neural network layers with rectified linear units between them and is capped by a SoftMax activation function to produce a normalized probability distribution denoting probabilistic state assignments as the final output.

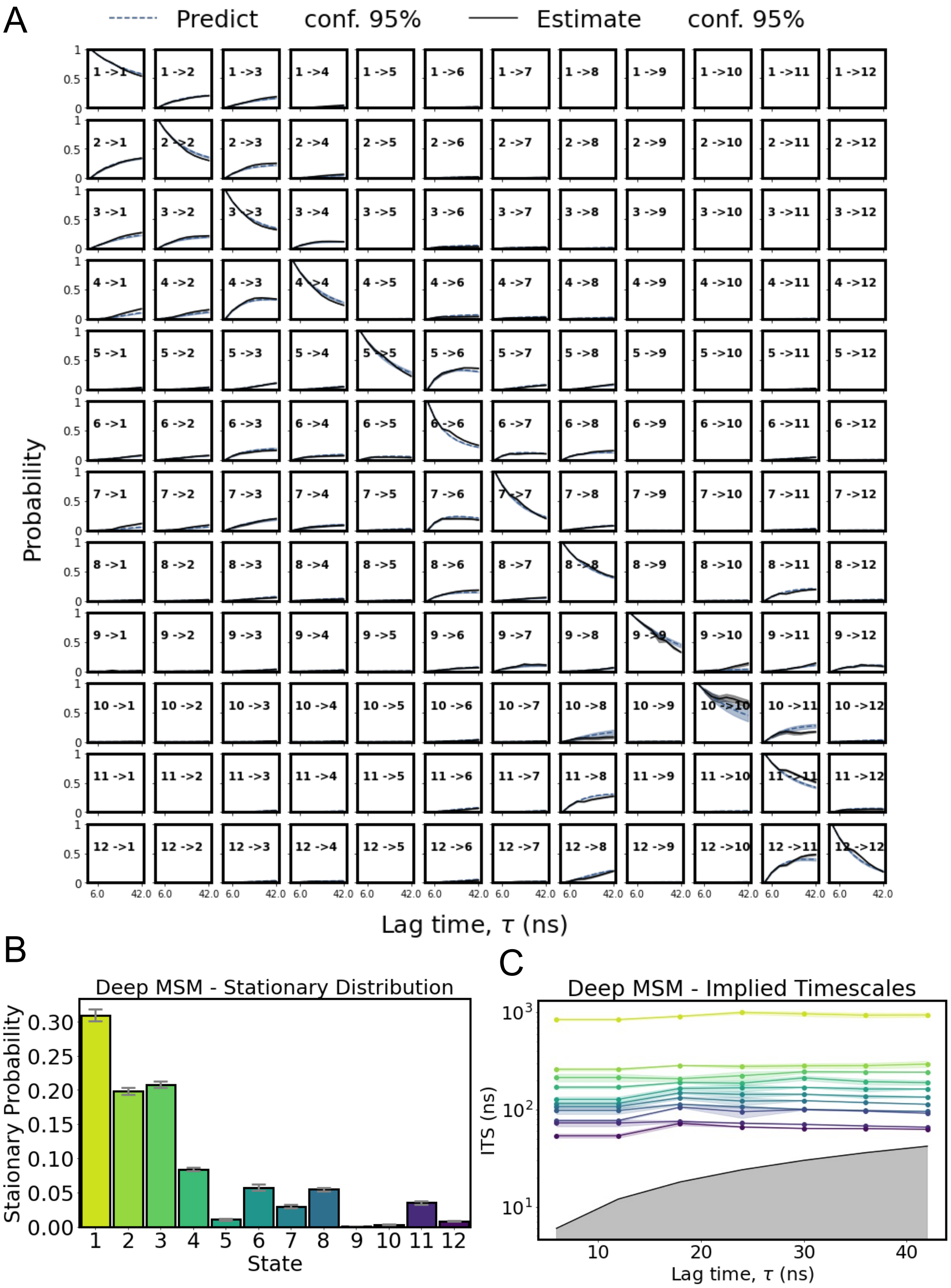


**Supplementary Figure 11. Deep MSM validation tests and stationary distribution. (A)** The Chapman-Kolmogrov (CK) test calculated for the deep MSM. The CK test evaluates the dependence of an MSMs predictions on the chosen lag time. The CK-test compares the evolution of transition probabilities for each state, i, to every other state, j, for integer multiples of the initial lag time of the model (6 ns). All quantities in the above plots were estimated individually for 30 transition matrices obtained from 30 separate optimization runs of the constrained VAMPnet and averaged (bootstrapped). Error bars were estimated by computing the 95% confidence of the bootstrapped means. Dashed blue lines ("Predict") represent the mean transition probabilities predicted by propagating deep MSM transition matrices estimated at fixed lag time (6 ns) and solid black lines ("Estimate") indicate the mean transition probabilities from transition matrices obtained from training only the constraint layers of the constrained VAMPnet at integer multiples of the initial lag time (6 ns). The blue and black shaded regions indicate 95% confidence intervals of the predicted and estimated transition probabilities, respectively. **(B)** The stationary distribution for each state of the deep MSM with error bars showing the deviation of the 95% confidence interval of the bootstrapped mean. **(C)** The log scaled implied timescales obtained from deep MSM transition matrices estimated at increasing lag times obtained by training only the constraint layers. The colored shaded regions show the deviation of the 95% confidence interval of the bootstrap mean. The solid black line and gray shaded region indicate time scales equal to or less than the lag time and represents the threshold for the fastest ITS that can be resolved by the model.

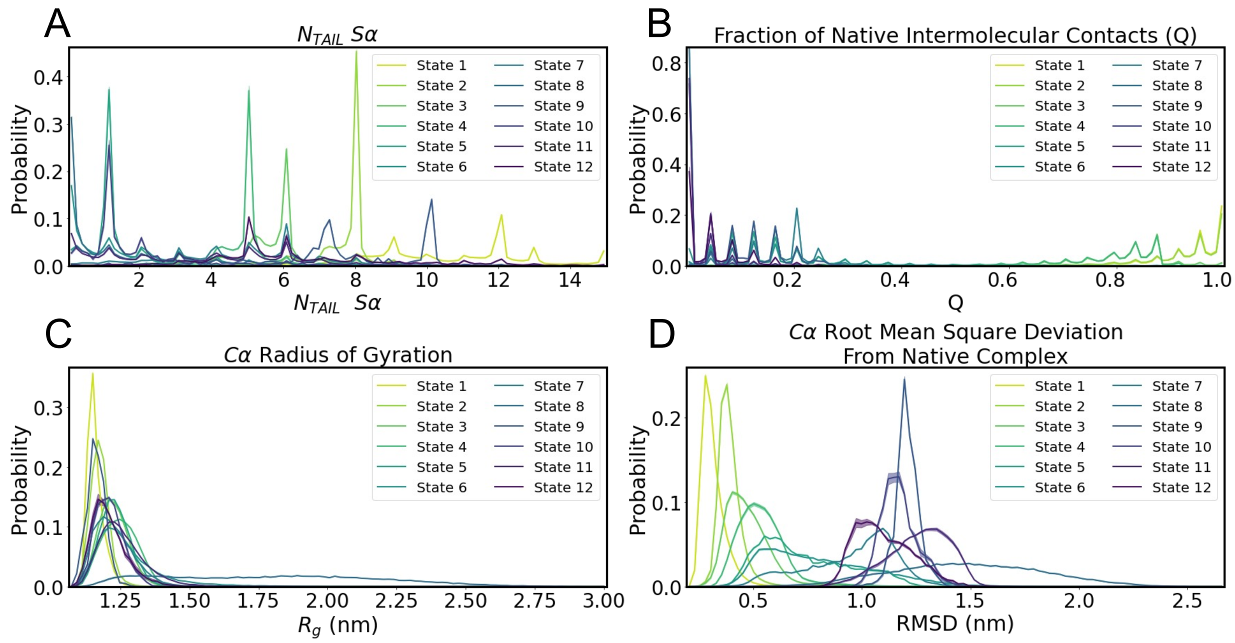


**Supplementary Figure 12. Structural properties of deep MSM states.** The discrete probability distributions of **(A)** $N_{TAIL}$ Sα (the number of ideal helical turns in $N_{TAIL}$), **(B)** the faction of native intermolecular contacts (*Q*), **(C)** the radius of gyration ($R_g$) computed using Cα atoms of $N_{TAIL}$ and XD and **(D)** the root mean squared deviation (RMSD) of $N_{TAIL}$ and XD Cα atoms from a solved X-ray crystal structure of the native complex (PDB 1T6O). The distribution of each statistic is computed separately for each state of the deep MSM, weighted by the probabalistic state assignments obtained from the constrained VAMPnet and averaged over the results of 30 independent optimization runs. The error bars in the distributions represent the 95% confidence interval of the bootstrap mean values.

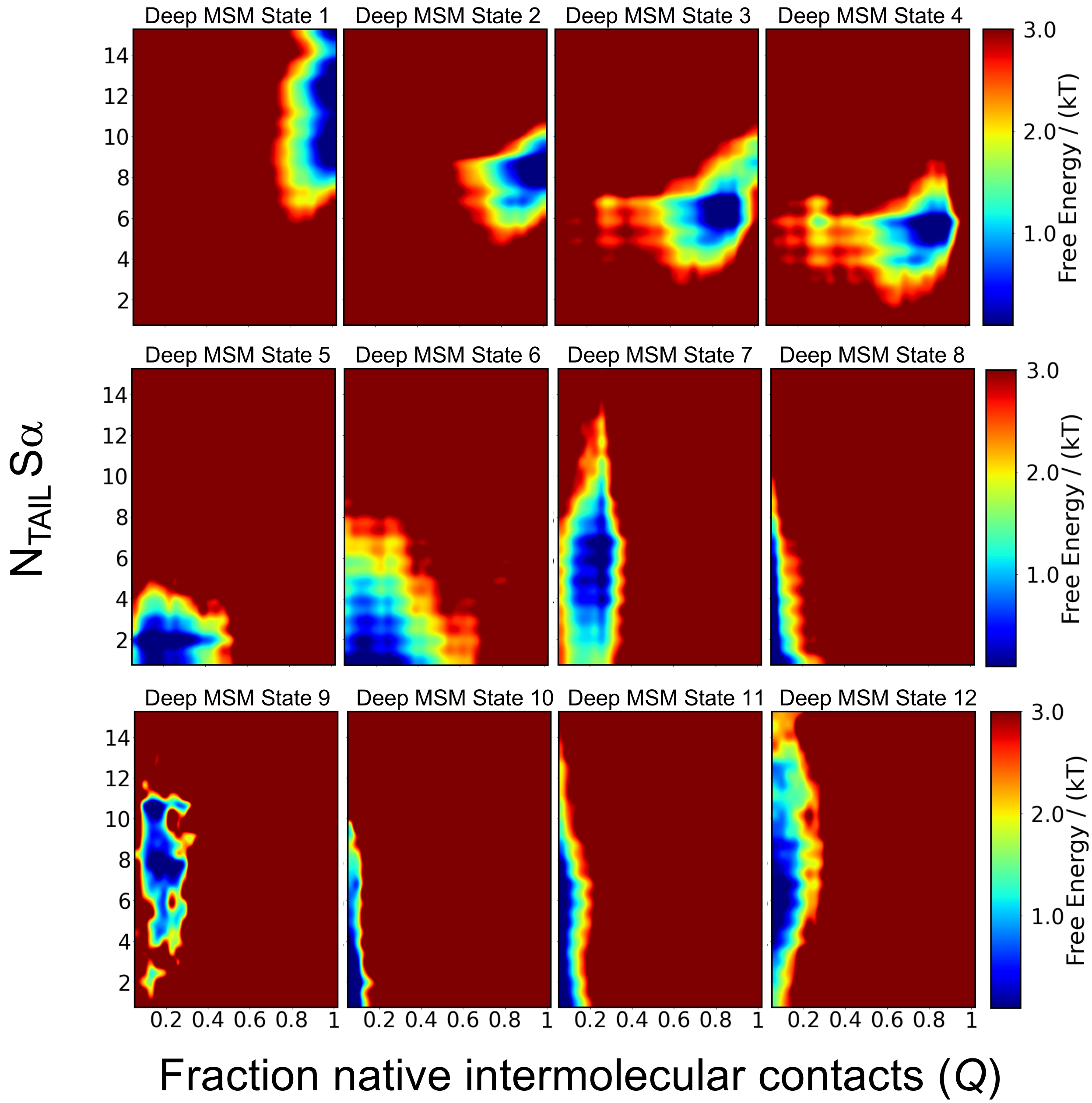


**Supplementary Figure 13. Free energy surfaces of the deep MSM conformational states as a function of $N_{TAIL}$ Sα and the fraction native intermolecular contacts (*Q*).** Free energy surfaces of the deep MSM states shown as a function of the fraction of native intermolecular contacts (*Q*) and $N_{TAIL}$ Sα (the number of ideal helical turns in $N_{TAIL}$) weighted by the state assignments obtained from the constrained VAMPnet and averaged (bootstrapped) over the results of 30 optimization runs.

| State | $\langle Q\rangle$ | $\langle Q\rangle_{SD}$ | $\langle N_{TAIL}\ S\alpha\rangle$ | $\langle N_{TAIL}\ S\alpha\rangle_{SD}$ | $\langle R_g\ (nm)\rangle$ | $\langle R_g\ (nm)\rangle_{SD}$ |
|---|---|---|---|---|---|---|
| 1 | 0.93 | 0.086 | 10.61 | 2.13 | 1.159 | 0.0279 |
| 2 | 0.91 | 0.11 | 7.66 | 0.92 | 1.175 | 0.0335 |
| 3 | 0.76 | 0.17 | 5.72 | 1.29 | 1.227 | 0.0541 |
| 4 | 0.73 | 0.19 | 4.75 | 1.01 | 1.249 | 0.0659 |
| 5 | 0.187 | 0.112 | 1.38 | 1.44 | 1.233 | 0.0542 |
| 6 | 0.193 | 0.158 | 1.95 | 1.96 | 1.276 | 0.129 |
| 7 | 0.183 | 0.09 | 5.05 | 2.32 | 1.229 | 0.0773 |
| 8 | 0.015 | 0.05 | 1.53 | 1.86 | 1.797 | 0.381 |
| 9 | 0.14 | 0.05 | 7.59 | 2.31 | 1.163 | 0.0327 |
| 10 | 0.013 | 0.022 | 1.89 | 1.69 | 1.210 | 0.0500 |
| 11 | 0.017 | 0.037 | 3.6 | 2.60 | 1.260 | 0.0970 |
| 12 | 0.052 | 0.058 | 6.52 | 2.70 | 1.209 | 0.0642 |

**Supplementary Table 2. Structural Properties of deep MSM States.** The bootstrap mean and standard deviation (SD) of the fraction of native intermolecular contacts *Q*, $N_{TAIL}$ Sα and the radius of gyration (nm) for each state of the deep MSM computed from the results of 30 separate optimization runs of the constrained VAMPnet. Here, we opt to display the pooled standard deviations computed from each trial to give a better estimate of the spread of each statistic, as opposed to the 95% confidence intervals of the bootstrap mean.

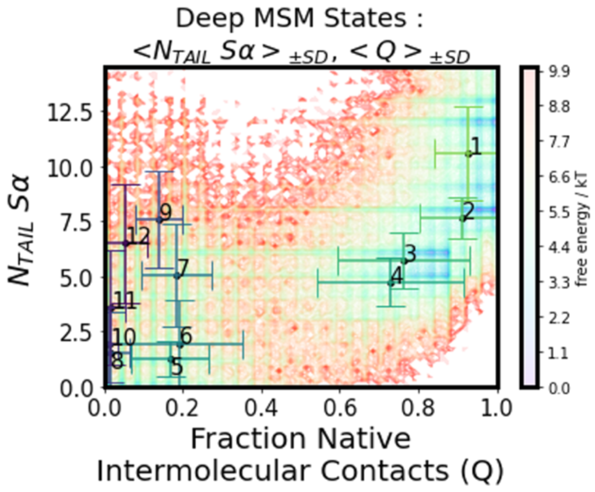


**Supplementary Figure 14. Comparison of properties of deep MSM states.** Free energy surface of $N_{TAIL}$:XD complex formation as a function of the α-helical folding order parameter Sα and the fraction of native intermolecular contacts, Q. The bootstrap mean of $N_{TAIL}$ Sα (<$N_{TAIL}$ Sα>) and Q (<Q>) are indicated for each deep MSM state. Error bars represent the standard deviation (SD) of each quantity, computed by aggregating the standard deviations obtained from the results of 30 optimization runs of the constrained VAMPnet.

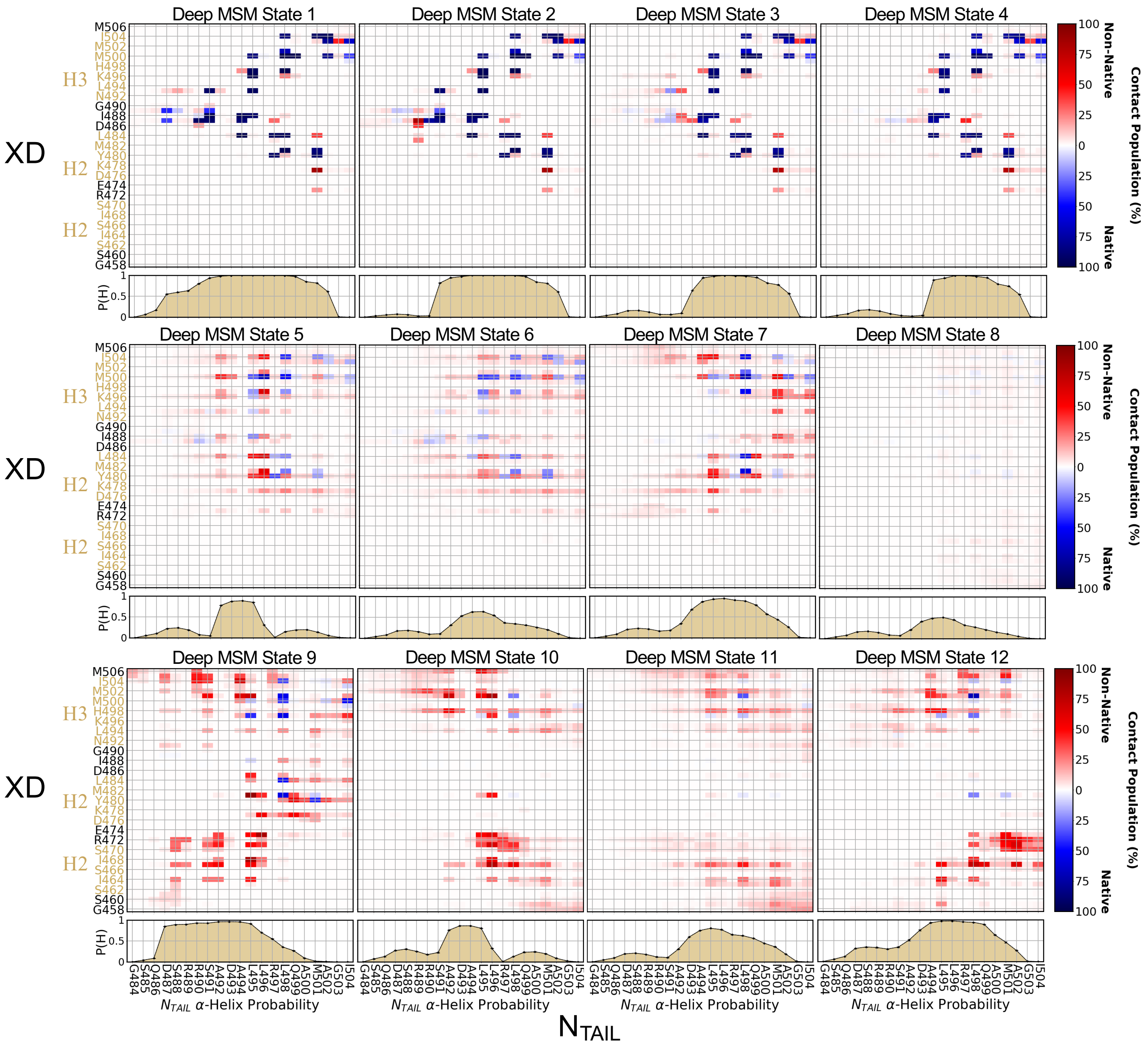


**Supplementary Figure 15. Intermolecular contact populations and $N_{TAIL}$ helical propensities of deep MSM states.** State averaged intermolecular $N_{TAIL}$:XD contact populations and $N_{TAIL}$ helical propensities for each deep MSM state. Intermolecular contacts were defined as occurring in all frames where the minimum distance between heavy atoms of two residues was less than 5.0 Å. Native intermolecular contact pairs are colored blue and non-native intermolecular contact pairs are colored red**.** Native contacts are defined as those present in the crystal structure (PDB 1T6O) using the same criteria. Helical propensities (P(H)) were calculated using DSSP. The mean contact populations and helix propensities of $N_{TAIL}$ are computed separately for each state of the deep MSM, weighted by the probabalistic state assignments obtained from the constrained VAMPnet and averaged (bootstrapped) over the results of 30 optimization runs.

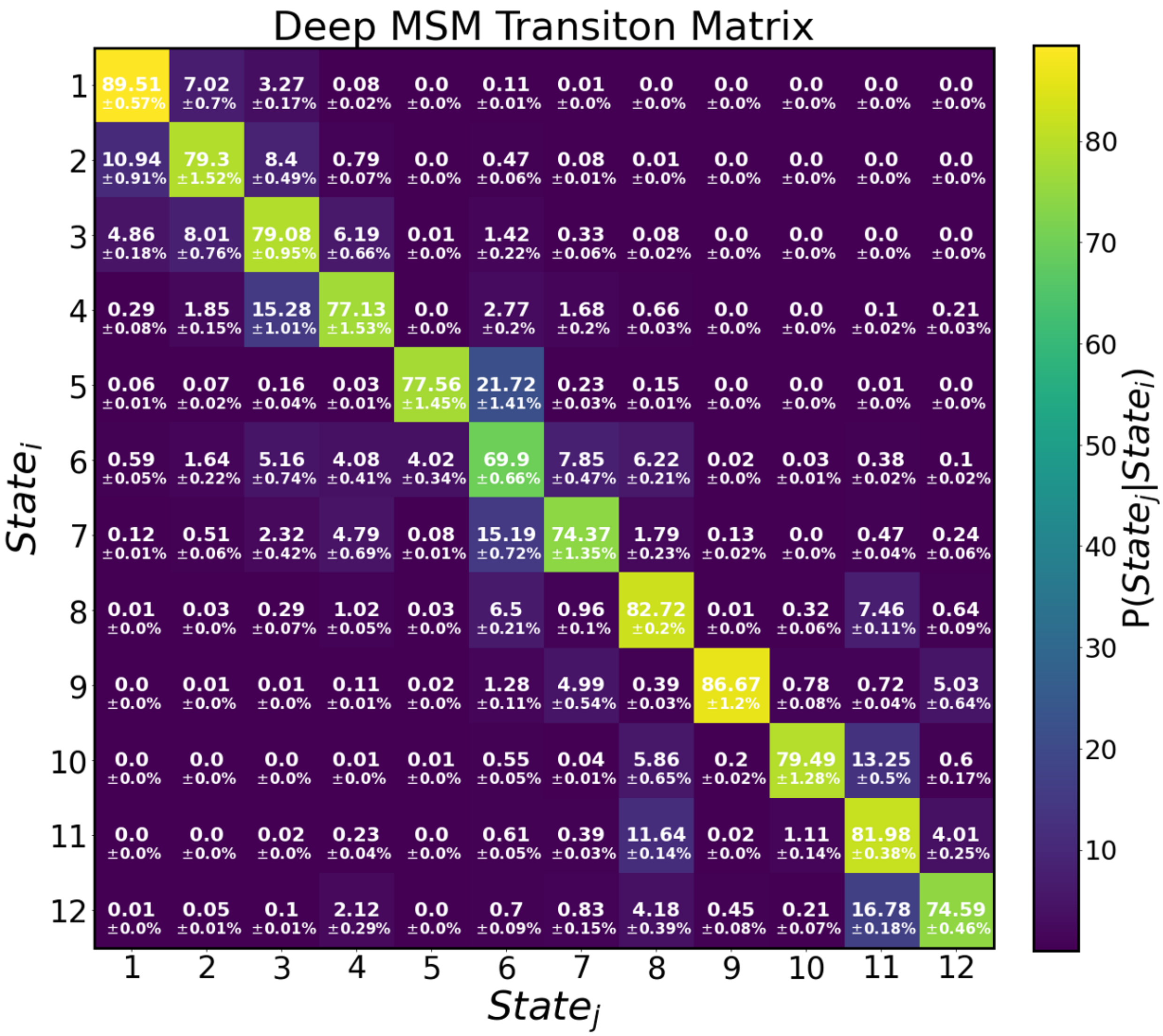


**Supplementary Figure 16. Deep MSM transition matrix.** The transition matrix for the deep MSM estimated at lag time τ = 6 ns. The transition matrix defines the conditional probability of transitioning from $state_i$ (at time t) to $state_j$ (at time t+ τ). The transition matrix shown here is the bootstrap mean of results obtained from 30 independent optimization trials of the constrained VAMPnet. Errors report the mean of the upper and lower deviations of the 95% confidence interval of the bootstrap mean.

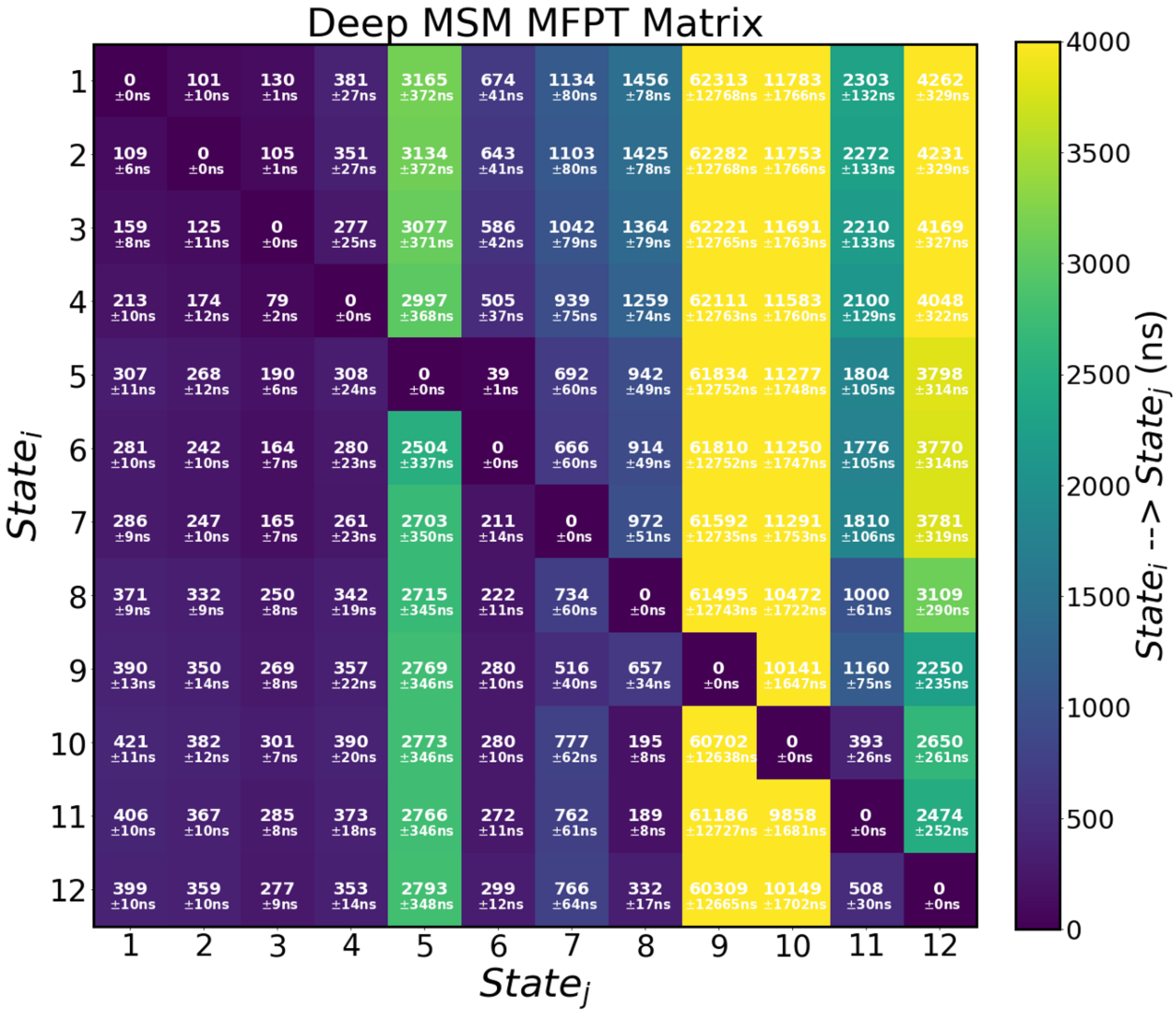


**Supplementary Figure 17. Deep MSM mean first passage time matrix.** The mean first passage time (MFPT) matrix shows the time duration it takes to transition between all Markov states of the deep MSM. We compute the MFPTs using the transition matrix and stationary probabilities of the deep MSM estimated at lag time $\tau = 6$ ns. The MFPT matrix shown here is the bootstrapped mean of results obtained from 30 independent optimization trials of the constrained VAMPnet. Errors report the mean of the upper and lower deviations of the 95% confidence interval of the mean. The mean first passages times estimated for states 9 and 10 have large errors in their estimation because of rarely being sampled in the original molecular dynamics' simulation. The color bar has a maximum value at 4000 ns, to provide visual contrast for the majority of MFPTs reported for well populated states. MFPT to the rarely populated states 9,10 are substantially larger than other transitions (MFPTs of ~10,000-63,000 ns) and are colored with the maximum value of the color bar.

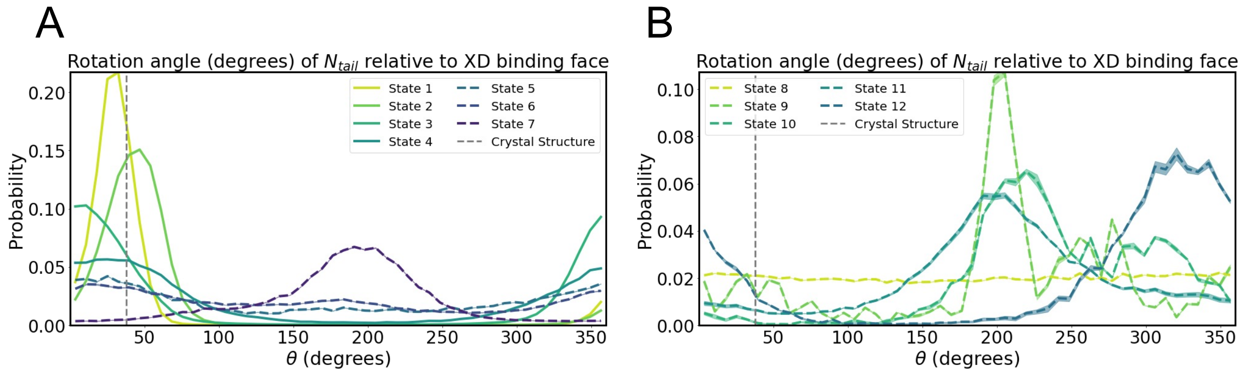


**Supplementary Figure 18. Orientation of $N_{TAIL}$ relative to the native binding interface of XD for each deep MSM state.** We quantify the orientation of $N_{TAIL}$ relative to the two helical bundles of XD (E475-L485 and N492-I504) forming the native binding interface as defined in Supplementary Appendix 1. The distribution of each statistic is computed separately for each state of the deep MSM, weighted by the probabalistic state assignments obtained from the constrained VAMPnet and bootstrapped over the results of 30 independent optimization runs. The error bars in the distributions represent the 95% confidence interval of the bootstrap mean values.

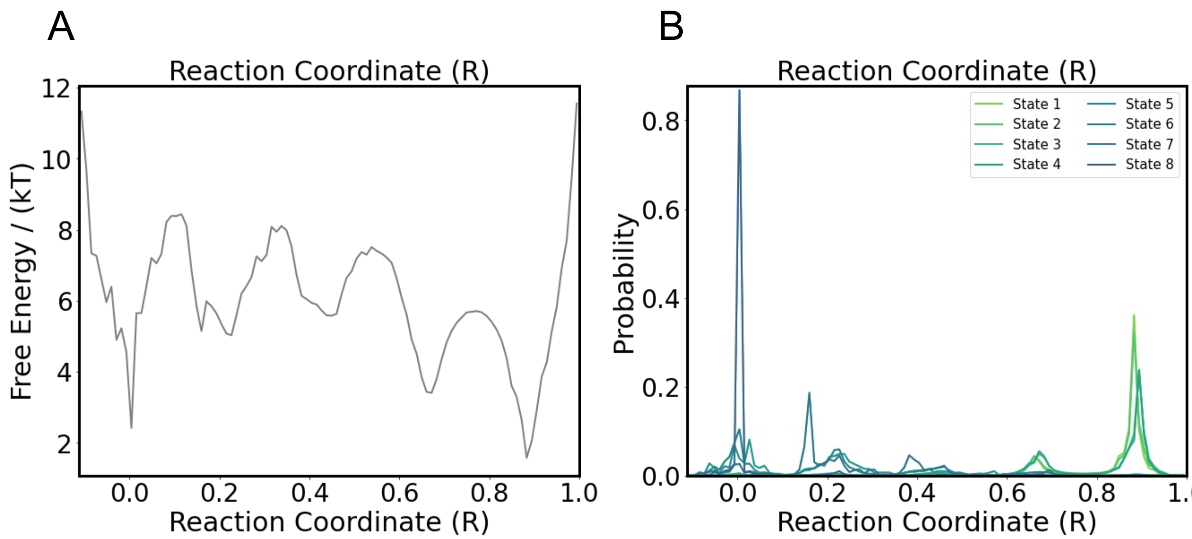


**Supplementary Figure 19. Apparent free-energy barriers observed with a previously defined 1D reaction coordinate for $N_{TAIL}$:XD folding-upon-binding are not kinetically meaningful**. **(A)** Free energy of the full $N_{TAIL}$:XD MD trajectory as function of the 1D folding-upon-binding reaction coordinated *"R"* previously defined by Robustelli et. al, *JACS*, 2020. **(B)** Probability distribution of the value of the reaction coordinate *R* for each kinetically distinct deep MSM state.

**Supplementary Appendix 1: Calculation of the orientation $N_{TAIL}$ relative to the helices that define the native XD binding groove.**

Here, all vectors are normalized to unit length at each step of the computation. We define the direction of propagation or orientation of a helical segment of a protein, $\vec{H}_{Propagation}$, as the mean of the nearest neighbor displacement vectors between alpha carbons computed from one fixed end of a segment to the other.

$$\vec{H}_{Propagation} = \frac{1}{N_{C\alpha} - 1} \sum_{i}^{N_{C\alpha}-1} \frac{\overrightarrow{C\alpha}_{i+1} - \overrightarrow{C\alpha}_i}{\left\| \overrightarrow{C\alpha}_{i+1} - \overrightarrow{C\alpha}_i \right\|} \quad (1)$$

For each frame of the trajectory, we compute the direction of propagation of $N_{TAIL}$, denoted as $\vec{H}_{NT}$, using the middle 19 residues of its sequence ($N_{TAIL}$:S485-G503) (we do not consider the terminal residues in the computation to reduce noise).

$$\vec{H}_{NT} = \frac{1}{19} \sum_{NT_{S485}}^{NT_{G503}} \frac{\overrightarrow{C\alpha}_{i+1} - \overrightarrow{C\alpha}_i}{\left\| \overrightarrow{C\alpha}_{i+1} - \overrightarrow{C\alpha}_i \right\|} \quad (2)$$

We compute the direction of propagation of the H2 (E475-L485) and H3 (N492-I504) helical bundles of XD that interface with $N_{TAIL}$ in the native folding upon binding process. We define the residues that comprise the helical bundles of XD based on the distribution of their sampled helical propensities.

$$\vec{H}_{XD_2} = \frac{1}{11} \sum_{XD_{E475}}^{XD_{L485}} \frac{\overrightarrow{C\alpha}_{i+1} - \overrightarrow{C\alpha}_i}{\left\| \overrightarrow{C\alpha}_{i+1} - \overrightarrow{C\alpha}_i \right\|}$$

$$\vec{H}_{XD_3} = \frac{1}{13} \sum_{XD_{N492}}^{XD_{I504}} \frac{\overrightarrow{C\alpha}_{i+1} - \overrightarrow{C\alpha}_i}{\left\| \overrightarrow{C\alpha}_{i+1} - \overrightarrow{C\alpha}_i \right\|} \quad (3)$$

We define a plane, $\sqcap_{XD}$, representing the interface of XD to which $N_{TAIL}$ binds. Using the cross product of $\vec{H}_{XD_2}$ and $\vec{H}_{XD_3}$, we define the normal vector ( $\vec{n}_{\sqcap_{XD}}$) of the plane that contains both $\vec{H}_{XD_2}$ and $\vec{H}_{XD_3}$. A plane is defined by a point on the plane and a vector normal to the plane, however, for our purposes we only require the normal vector.

$$\vec{n}_{\sqcap_{XD}} = \vec{H}_{XD_2} \times \vec{H}_{XD_3} \quad (4)$$

The mean of the $\vec{H}_{XD_2}$ and $\vec{H}_{XD_3}$ orientation vectors is computed to obtain the orientation of the XD binding interface.

$$\tilde{\vec{H}}_{XD} = \frac{\vec{H}_{XD_2} + \vec{H}_{XD_3}}{2} \quad (5)$$

Note that as $\vec{H}_{XD_2}$ and $\vec{H}_{XD_3}$ both sit in the $\sqcap_{XD}$ plane, so does their mean or any linear combination of them. The NTAIL orientation vector is projected unto the plane of the XD binding interface.

$$Proj_{\sqcap_{XD}}\vec{H}_{NT} = \vec{H}_{NT} - \vec{H}_{NT} \cdot \ \vec{n}_{\sqcap_{XD}} \qquad (6)$$

As a result of this projection and normalization of all vectors at each step of the computation, the deviation between $\tilde{\vec{H}}_{XD}$ and $\vec{H}_{NT}$ can be fully captured by a single angular polar coordinate, $\theta_{NT:XD}$.We compute the signed sin and cos of $\theta_{NT:XD}$ and use them as arguments to the arctan2 function in to recover the full angle, defined between 0 and 360 degrees.

$$\begin{gathered}\cos\left(\theta_{NT:XD}\right) = \tilde{\vec{H}}_{XD} \cdot Proj_{\sqcap_{XD}}\vec{H}_{NT} \\ \sin(\theta_{NT:XD}) = (\tilde{\vec{H}}_{XD} \times Proj_{\sqcap_{XD}}\vec{H}_{NT}) \cdot \vec{n}_{\sqcap_{XD}} \\ \theta_{NT:XD} = \text{atan2}[\sin(\theta_{NT:XD}), \cos\left(\theta_{NT:XD}\right)]\end{gathered} \qquad (7)$$
